# Social License Pressure and Public Support for the Lethal Control of Dingoes in Australia

**DOI:** 10.1101/2025.01.20.633982

**Authors:** A I Pearson, M L Cobb

## Abstract

The lethal control of dingoes to protect livestock has been publicly funded since Australia was colonised. Although lethal dingo control had social license to operate (i.e., public support) from colonial settlers, there have been limited examinations into contemporary Australian opinions. This study explored predictive factors for Australian public support for lethal dingo control. It was hypothesised that higher social license pressure (perceptions of low procedural fairness, trust, and confidence in the government’s dingo management policies), positive attitudes toward Australian wildlife, and dog ownership would predict less support for lethal dingo control. Additionally, attitude towards Australian wildlife was predicted to mediate the relationship between social license pressure and support for lethal dingo control. Australian adults (*N* = 990, aged 18 – 83, 52% women) voluntarily completed an online survey, which included the Attitudes to Australian Wildlife scale and novel items. Results supported all hypotheses (*p* < .001). More Australians opposed lethal dingo control than supported it. Social license pressure, attitude towards Australian wildlife, and dog ownership significantly predicted support for lethal dingo control as hypothesised. Attitude towards Australian wildlife mediated the relationship between social license pressure and support for lethal dingo control with a large effect size (*f^2^* = 1.73). This study provides evidence of tenuous community support for the lethal control of dingoes. Given that a loss of social license would challenge Australia’s longstanding livestock practices, our findings are particularly useful for the proactive development of dingo management policies that the Australian public will support.

**Highlights:**

- Support for lethal dingo control was polarised, but more Australians were opposed
- Unfavourable public perception of dingo policies; increasing social license pressure
- Dog owners demonstrated less support for lethal dingo control than non-dog owners
- Wildlife attitudes mediated social license and support for dingo control relationship
- Social license to operate for lethal dingo control is tenuous in Australia

## 1. Introduction

Social license pressure is an increasingly visible process in economically advanced democratic countries. It refers to the informal yet tangible influence of societal expectations on the activities of organisations and institutions beyond their legal obligations (Graafland & Smid, 2017). Although most evident in extractive natural resource industries (Moffat et al., 2016), social license pressure has led to changes in longstanding animal-use practices in Australia, including the lethal control of wildlife (Hampton et al., 2020). Public support for wildlife lethal control varies across many factors. These include the wildlife species, the wildlife’s perceived charisma, whether the wildlife is consumed or commercially sold, political context, media coverage (type, availability, and valence), attitudes and emotions toward wildlife, and the values demonstrated by a population at a given time (Arbieu et al., 2019; Drijfhout et al., 2020; van Eeden et al., 2017).

Dingoes are one wildlife species that have been targeted by costly, yet largely ineffective, publicly funded lethal control efforts to protect livestock since Australia was colonised (Corbett, 2001). The cost of dingo management in Australia is large. It has been estimated at an annual cost of AU$13 - $27 million (Brink et al., 2019; Smith & Appleby, 2018; Wool Producers Australia, 2014). Although the Australian population has changed considerably in the last two centuries, lethal dingo control has remained a publicly funded policy without investigation as to whether this aligns with public opinion (van Eeden et al., 2021). It is therefore important to understand the Australian public’s support for lethal dingo control, and how this intersects with social license pressure and attitude towards Australian wildlife. Such findings will further extend social license theory into the field of wildlife management. It can also add to human-animal intersection literature, which includes the legal rights of non-human animals (Vink, 2020). Further, this research serves to inform and shape government wildlife policy and practices, and the development of attitudinal interventions, and educational material for stakeholders, including the public.

### 1.1. Dingoes

Dingoes are found across mainland Australia and have been controversial since they were first mentioned in written European records, particularly their naming. Taxonomic debate includes *Canis lupus dingo*, *Canis familiaris* (breed Dingo), *Canis dingo*; while they are classified as free-ranging canids of the Canidae family which includes coyotes and wolves (Smith, 2015), the term used for dingoes is inconsistent. The term of “dingo” is used in literature focussed on conservation and coexistence, and that originates in Australian states (e.g., Northern Territory, Western Australia) where dingo purity has typically been considered higher (Kreplins et al., 2019; Smith & Appleby, 2018). Although the classification of a “pure” dingo is debated (Crowther et al., 2014; Smith & Appleby, 2015), newer DNA testing methods have demonstrated that most Australian wild canid populations should be classified as dingoes (Cairns et al., 2023). This holds even in south-eastern Australia where studies based on older DNA testing methods indicated that large-scale hybridisation with domestic dogs had occurred (Cairns et al., 2020; Stephens et al., 2015).

The term of “wild dog”, however, is used elsewhere. For example, by Australian print media (Degeling et al., 2021), state governments (Letnic, 2012), and livestock-related organisations (Australian Wool Innovation Ltd, 2020). This term also includes feral domestic dogs, unowned stray dogs, unrestrained free-ranging dogs, and dog-dingo hybrids (Kreplins et al., 2019). The wild dog term is also used in literature that originates in Australian east-coast states where studies based on older DNA testing methods suggested that dingo purity was limited (Cairns et al., 2023; Kreplins et al., 2019). What term is used to describe dingoes has implications for their treatment.

Dingoes are treated differently depending on which term is used. Government and industry documents differentiate between the “control” of wild dogs (e.g., Australian Wool Innovation Ltd, 2020), and the “protection” of dingoes in places like national parks (e.g., Queensland Government, 2023). This is challenging because of the visual similarity of domestic dogs and dingoes (Smith, 2015), and because the definition of wild dogs in these documents includes dingoes. Notably, Indigenous Australians (Costello et al., 2021), and the general Australian public consider dingoes to be separate from wild dogs. For example, 81% of a representative national sample (*N* = 811) recruited in proportion to census data were unaware that the two terms encompassed the same animal (van Eeden et al., 2021). Since the current study focussed on assessing the Australian public’s perspective, the term dingo was adopted.

### 1.2. Dingo Control

Since European colonisation, dingoes have been controlled by lethal and non-lethal means. Lethal means have been dominant. These include killing by shooting, trapping, and denning, with the most common method being ground and aerial baiting with strychnine or sodium monofluoroacetate (compound 1080) (Binks et al., 2015). Non-lethal control has typically been exclusion fencing, most notably the dingo barrier fence that covers three Australian states (Woodford, 2003). Historical data indicates that over 1.5 million dingo scalps (from trapping) were presented for bounty payment between 1885 and 1995 in Queensland alone (Allen & Sparkes, 2001). However, there are methodological difficulties (e.g., most poisoned dingoes are not recovered) which suggest that dingo death counts are higher than those reported over the last two centuries.

The longstanding rationale for dingo control has been that dingo populations must be reduced to decrease the loss of livestock from predation. The loss of livestock from dingoes has been estimated at AU$89 million annually (average cost in 2013-2014 dollar terms; McLeod, 2016). Psychological trauma is also reported by afflicted individuals. For example, Ecker et al. (2017) found that stress impact scores for landholders experiencing livestock predation attributed to dingoes were comparable to the scores of male Vietnam veterans in the general Australian community experiencing a range of traumatic stress symptoms (although not a post-traumatic stress disorder diagnosis). While these results must be interpreted cautiously due to it being a pilot study, small sample size, and lack of researcher independence,^1^ it nonetheless highlights that there are (non-economic) human costs for livestock producers and their communities from livestock loss.

There is limited peer-reviewed evidence that shows reducing dingo or top-predator numbers protects livestock, let alone directly reduces livestock loss. Research has shown that culling dingoes has a negligible effect on cattle numbers (Allen, 2015; Fleming et al., 2015), and has even worsened cattle losses (Allen, 2014; Campbell et al., 2019). The positive correlation between predator culling and livestock loss, such that livestock losses increase (i.e., worsens) the year after a predator cull, has also been seen in wolf-livestock studies in Spain (Fernández-Gil et al., 2016), and the United States (Wielgus & Peebles, 2014). Longitudinal sheep studies in the United States have shown that coyote baiting did not change sheep losses (1948 – 1955; Wagner, 1972), or made only a minor independent contribution (6%; 1950 – 1970; Berger, 2006). This echoes reviews that found mixed evidence of the effectiveness of lethal carnivore control to protect livestock (van Eeden, Crowther, et al., 2018; van Eeden, Eklund, et al., 2018). Together, the limited and contradictory evidence base suggests that factors outside reducing predator populations are important to reduce livestock loss.

While the underlying mechanisms of the predator-livestock loss relationship are debated in the literature, research has demonstrated associations between indiscriminate top-predator culling and the fracturing of canid pack stability. Naïve and non-resident predators, such as dingoes (Allen, 2015) and wolves (Shivik et al., 2003) appear more likely to target livestock than stable predator packs. When dingo pack hierarchies are destabilised by culls, it can paradoxically lead to increased dingo numbers through compensatory dingo breeding by other females if the former breeding female is killed (Glen et al., 2007). Dingo culling can also lead to the loss of livestock and their feed because other predators, mesopredators, and herbivores are no longer constrained by stable dingo packs that defend their territory (Johnson et al., 2014; O’Neill et al., 2017). Furthermore, livestock loss may worsen irrespective of reduced predator numbers, because particular individuals or pairs of dingoes (Allen & Fleming, 2004) and coyotes (Sacks et al., 1999) have been associated with high predation rates on their own. Given that the long and expensive interventions to reduce dingo numbers have not led to reduced livestock loss, the rationale for lethal dingo control policies must be questioned.

The endurance of the rationale that lethal dingo control reduces livestock loss despite a lack of empirical support sits within an interplay of factors. Lethal dingo management practices can provide a sense of control for livestock producers, alongside a social identity of “good neighbour, good farmer” as defined by the traditions of colonial Australia (Boronyak et al., 2023). Agricultural interest groups are also politically powerful in Australia (van Eeden et al., 2017), which reinforces government bias for lethal control. Government support prioritises free or subsidised bait, bounties, and training in lethal control of dingoes for livestock producers (Hillier, 2017). This then reinforces the perception of lethal control as a normal management strategy in rural communities (Treves & Naughton-Treves, 2005; Van Eeden et al., 2019). Such factors and their interactions serve to reinforce lethal dingo control as the mainstay policy to protect livestock in Australia. Whether this policy is supported depends on the time period, and which members of the Australian public are being consulted.

### 1.4. Australian Public Support for Lethal Dingo Control

Historically, lethal dingo control was publicly supported by colonial settlers who established agriculture as the backbone of Australia’s economy and national identity. Newspapers and other written material described dingoes as menaces and pests, reporting strong support for lethal dingo control amongst British and Irish settlers (Breckwoldt, 1988; Sidney, 1854). This contrasted with the long tradition of harmonious coexistence with dingoes practised by Indigenous Australians, whose opinions have rarely been sought (Costello et al., 2021). The post-colonisation land use choices impacted dingo numbers and behaviour by introducing anthropogenic food sources (e.g., sheep and cattle) and changing water sources (Corbett, 2001; Smith et al., 2019). Research has suggested that the longevity and severity of dingo-livestock conflict in western New South Wales compared to other areas can be partially attributed to the area’s discovery by sheep (rather than cattle) graziers (Newsome, 2001). This is because sheep exhibit flight and mobbing behaviour that favour successful predation by dingoes compared to cattle (Allen & Fleming, 2004). Australian public support for lethal dingo control therefore sits within a broader historical and cultural context that spans more than two centuries.

The enduring designation of dingoes as pests continues to underlie Australia’s dingo control policies. Most Australian jurisdictions today include dingoes in their wild dog classification (see summary by Australian Wool Innovation Ltd, 2020). This requires landholders to legally control them as biosecurity threats (Queensland Government, 2021). This aligns with evidence that people’s attitudes differ depending on the context of an animal. Taylor and Signal (2009) found that whether an animal was considered profitable (e.g., livestock), useful (e.g., livestock guardian animal), a pest (e.g., predator), or a pet (e.g., companion animal), impacted people’s attitude towards them. Whether contemporary Australians continue to support the lethal control of dingoes, more than two centuries after colonisation, must be considered in the context of modern-day Australia.

### 1.5. Contemporary Australia

There are many changes between colonial and contemporary Australia. Agriculture was colonial Australia’s biggest industry. It accounted for 19.4% of Australia’s gross domestic product in 1901, compared to only 2.7% in 2022 (Australian Bureau of Agricultural and Resource Economics and Sciences [ABARES], 2024). In 1911, Australians were relatively evenly spread between urban (58%) and rural (42%) areas. However, by 2016, 90% of Australians lived in urbanised areas and only 10% lived in rural areas (Australian Bureau of Statistics [ABS], 2019). Despite this, livestock agriculture remains widespread. More than half (54%) of the Australian continent is used for livestock grazing (ABARES, 2016). Of these farm businesses, 38% raise cattle or sheep, or both (ABARES, 2023). Further, it is estimated that more than AU$30 million is spent annually by governments and landowners on dingo control (Brink et al., 2019). Together, this suggests that dingo-livestock conflict remains a topical concern in Australia.

### 1.6. Contemporary Opinions on Dingoes

There is preliminary evidence that Australians view dingoes differently now, than they did in the past. In an Australian sample recruited proportionally to census data (*N* = 811), van Eeden et al. (2021) found that 53% of participants did not consider the dingo a pest, and 65% were unaware that dingoes were legally controlled by lethal means. Most respondents (81%) believed that wild dogs were different to dingoes, and held positive attitudes towards dingoes (although negative attitudes towards wild dogs). They disapproved of all lethal control methods for dingoes, however supported non-lethal strategies (e.g., exclusion fencing). Since attitudes differ as a function of time and social influences (Heberlein, 2012; Jackman & Rutberg, 2015), these results must be interpreted in line with the characteristics of contemporary Australians.

Contemporary Australians generally live away from dingoes and value animals. Most Australians live in urbanised areas (90%; ABS, 2019) with limited first-hand knowledge of rural animal practices (Coleman, 2018). Cross-cultural and longitudinal studies have demonstrated that an urban upbringing, not identifying with rural stakeholder groups (e.g., hunters or agricultural producers), living at greater geographical distance from wildlife, and being less familiar with wildlife, are associated with more positive attitudes toward wildlife and their conversation (Karlsson & Sjöström, 2007; Landon et al., 2019; Vaske et al., 2022; Williams et al., 2002). Australians also report high pet ownership (69%), predominantly of dogs (48%; Animal Medicine Australia, 2022), and value animal welfare (Chen, 2016; Coleman, 2018). Their moral objection to animal suffering has been fuelled by media exposés and social media activism (Hampton et al., 2020), and resulted in public protest against previously accepted animal-use industries in Australia. Examples include greyhound racing (Markwell et al., 2017) and live cattle exports (Schoenmaker & Alexander, 2012). This shift towards social disapproval of using animals for human benefit when animal welfare is compromised suggests that social license pressure may be increasing in Australia.

### 1.7. Social License Pressure

As part of the social license to operate concept that originated in the mining sector (Cooney, 2017), social license pressure is the control that community expectations can exert on organisations and institutions that use resources with a perceived public value (Graafland & Smid, 2017; Lynch-Wood & Williamson, 2007). Although intangible (i.e., cannot be granted legally or politically; Morrison, 2014), social license pressure has demonstrated real impacts on wildlife management practices. For example, strong public disapproval stopped an intended cull of Australian feral horses (i.e., brumbies) in 2000 (Nimmo et al., 2007). Conversely, a successful cull of Australian feral camels was undertaken with community support, following proactive consultation with stakeholders which included animal welfare groups (Hart & Bubb, 2016). Social license pressure therefore affects wildlife management; however, models have only been validated for extractive sectors.

### 1.8. Measuring Social License Pressure

After consulting existing validated social license tools, the current study measured social license pressure through the elements of procedural fairness, trust, and confidence. The most recent model validated for mining in Australia, China and Chile (Zhang et al., 2015) found four social license components: distributional fairness, confidence in governance, procedural fairness, and trust. The current study excluded distributional fairness (i.e., equitable distribution of mining’s economic benefits), and confidence in governance (i.e., government legislation holds the mining industry accountable) as these were not relevant to government lethal wildlife control policies. Procedural fairness and trust were included in the current study. However, because Zhang et al. (2015) did not define trust, and wildlife management studies often conflate trust and confidence (e.g., Engel et al., 2016), the broader literature was consulted.

The current study’s measurement of trust and confidence as facets of social license pressure was informed by social and institutional risk research. Here trust is based on integrity, and confidence is based on competency (Siegrist, 2021). This has been supported by factor analyses and experimental manipulations that found trust and confidence uniquely predict social cooperation and acceptance of collective decisions (Siegrist et al., 2012; Twyman et al., 2008). Additionally, a principal components analysis found that acceptance by hunters of decisions taken by a United States wildlife authority (which may also be referred to as wildlife department, agency, etc.) was underpinned by perceptions of the authority’s integrity, competency, and procedural fairness (Schroeder et al., 2017). Together, this suggests that procedural fairness, trust, and confidence are appropriate facets to measure as part of social license pressure in the current study.

#### 1.8.1. Procedural Fairness

Following Tyler (2015), the current study defined procedural fairness as the public’s perception that they were reasonably included and respectfully treated during dingo management decision-making processes. Studies have found positive associations between procedural fairness and acceptance of socially polarising or personally unfavourable outcomes. These include the location of nuclear power plants (Besley, 2010), genetically modified crops (Siegrist et al., 2012), and mining activities (Zhang et al., 2015). Research suggests that acceptance occurs because procedural fairness confers valuable social and relational benefits. Tyler (2015) found that people’s social rights, status, and identity are affirmed when they perceive their opinions are duly received by authorities on a “level playing field”. Together, this indicates that perceptions of procedural fairness are important in understanding support for lethal dingo control.

The limited evidence base investigating procedural fairness and public support for wildlife management practices suggested a positive relationship, whereby perceptions of higher procedural fairness correlate with heightened public support. Research on moose reintroduction in the United States found that when procedural fairness was rated higher, the public was more satisfied with the decision-making process and the wildlife authority (Lauber & Knuth, 1999). Similarly, Sjölander-Lindqvist et al. (2015) found that acceptance of management decisions for large carnivores (e.g., wolf, lynx, brown bear) in Sweden was enhanced when local representation was considered fair and comprehensive. Together, this suggests that procedural fairness (as a component of social license pressure) and public support for lethal dingo control should have a positive relationship. Specifically, higher procedural fairness (indicative of lower social license pressure) would be correlated with greater support for lethal dingo control.

#### 1.8.2. Trust

Following Poppo and Schepker (2010), the current study defined trust as the public’s perception that the government was acting and communicating with integrity (i.e., what is “right”) about dingo management. Research has found that when public knowledge about the risks of a socially contentious activity is low, social and institutional trust has positively predicted acceptance. Examples include support for nuclear power (Vainio et al., 2017), genetically modified crops (Siegrist et al., 2012), and mining (Moffat & Zhang, 2014). In such scenarios, trust acts as a heuristic, allowing people to continue operating in complex situations without effortfully acquiring sufficient knowledge to make deliberate assessments about acceptability, favourability, and risks (Siegrist, 2021). This suggests that whether the government’s dingo management practices are perceived to be integrous (i.e., ethical and honest) would affect support for lethal dingo control.

Although there is limited precedent for our definition of trust, research suggests that trust would be positively associated with support for wildlife management. Trust in the wildlife authority positively predicted acceptance by anglers of the United States fisheries’ advice and non-specified future management decisions (Schroeder & Fulton, 2017). Similarly, higher ratings of trust in a wildlife authority by residents, hunters and anglers positively predicted greater satisfaction with the authority’s conservation efforts (Connelly et al., 2022). Together, the research indicates that trust (as a component of social license pressure) and public support for lethal dingo control would be positively related. Specifically, higher trust (indicative of lower social license pressure) would be correlated with greater support for lethal dingo control.

#### 1.8.3. Confidence

Following Earle et al. (2007), the current study defined confidence as the public’s expectation that the government was sufficiently knowledgeable and experienced (i.e., competent) to manage dingoes. Such confidence has been positively correlated with acceptance of socially controversial decisions. Examples include nuclear power plant expansion (Besley, 2010), carbon dioxide capture and storage technology implementation (Terwel et al., 2009), and the residential locating of electromagnetic fields (Siegrist et al., 2003). In a similar mechanism to trust, confidence allows people to believe that an institution has the appropriate expertise to manage a situation on their behalf (Siegrist, 2021). This suggests that whether the public perceived the government to be competent in dingo management would influence their support for lethal dingo control.

Research has shown that confidence in wildlife authorities is positively associated with public support for wildlife management practices. This has been demonstrated in research about bears, cougars, and wolves (Ghasemi et al., 2021). Even amongst hunters, those who lacked confidence in the wildlife agency’s moose data were nearly five times more likely to consider increased quotas on antlerless moose harvesting as unacceptable (Brinkman, 2018). Together, this proposes that confidence (as a component of social license pressure) and public support for lethal dingo control would be positively related. Specifically, higher confidence (indicative of lower social license pressure) would be correlated with greater support for lethal dingo control.

In sum, the attitudes held by members of the public matter when it comes to community support for the management of wildlife. The reviewed research has suggested that attitudes consistent with lower social license pressure, as measured by more positive (i.e., higher) perceptions of procedural fairness, trust, and confidence about the government’s management of dingoes, would likely be associated with higher support for lethal dingo control (see Figure 1, pathway *c*). Additionally, whether people demonstrate positive or negative attitudes toward wildlife changes their support for wildlife management practices. For example, Drijfhout et al. (2020) found that lethal culling was acceptable for kangaroos but not for koalas in a sample of Australian adults (*N* = 1,148) recruited proportionally to census data. The authors suggested this difference may be related to participants holding more negative attitudes about kangaroos because they threaten human life in ways (e.g., fatal traffic accidents, crop damage) that koalas do not. This suggests that in addition to social license pressure, understanding the valence of attitudes held toward Australian wildlife may assist in predicting support for lethal dingo control.

**Figure 1.**
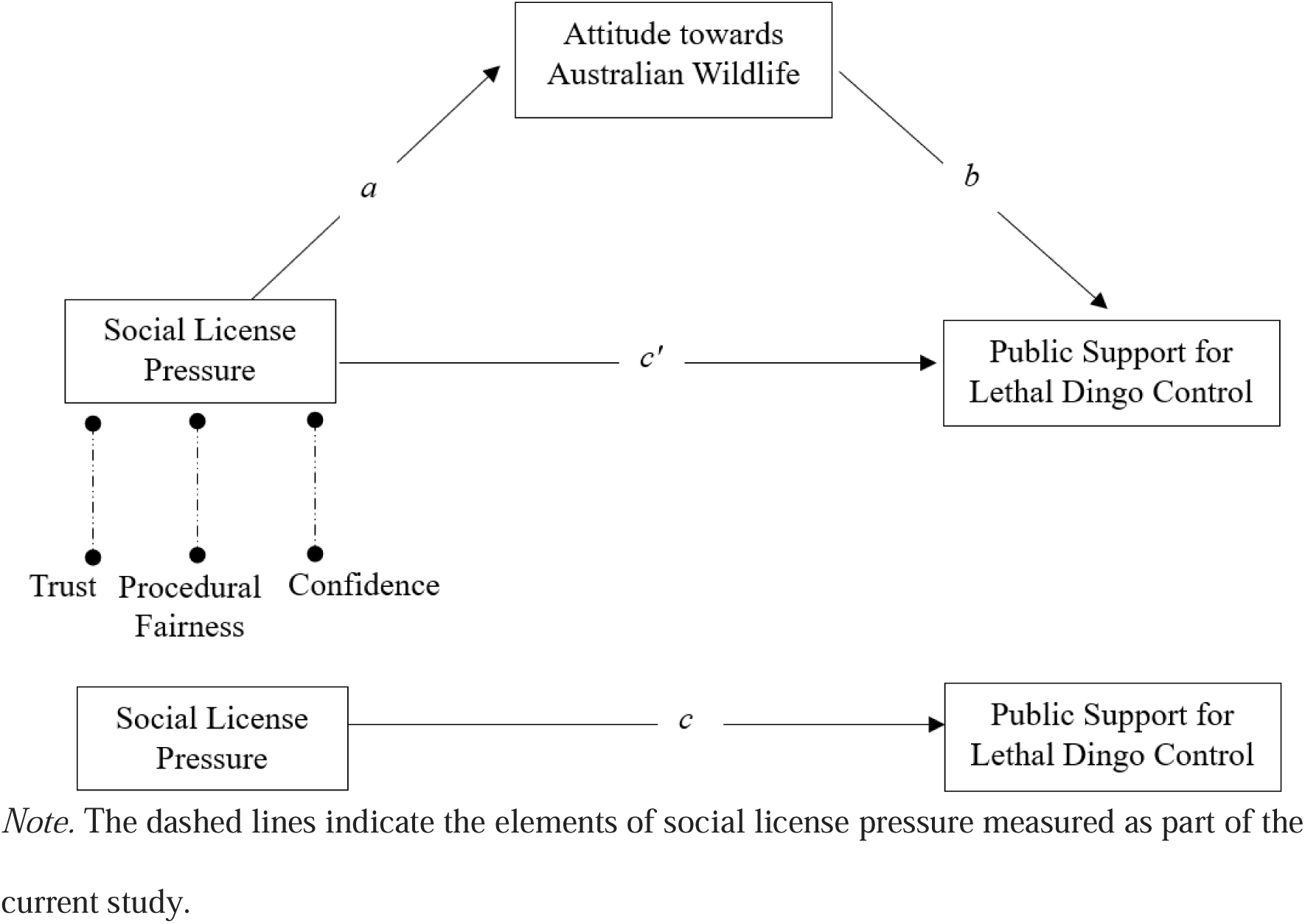
Proposed Relationships Between Social License Pressure and Public Support for Lethal Dingo Control as Mediated by Attitude towards Australian Wildlife

### 1.9. Attitude towards Australian Wildlife

While wildlife attitudinal investigations are limited in Australia (Boulet et al., 2021), Australians appear to have positive attitudes toward native wildlife and disapprove of lethal wildlife control. Compared to university students from six other countries, Jacobs et al. (2022) found that Australian students demonstrated the highest scores on mutualistic wildlife values orientation (i.e., wildlife should be extended care and rights like humans), and the most negative attitudes toward using wildlife for human benefit (e.g., hunting). Although caution is warranted because only students from one Australian university were surveyed, the comparative strength of Australian opposition to harming wildlife may be related to Australia’s high rate of wildlife extinction (Johnson, 2006), and the uniqueness of Australian ecosystems where more than 80% of plant and animal species are endemic (Alacs & Georges, 2008). A negative relationship between attitude towards wildlife and public support for lethal wildlife control, such that a more positive attitude towards the wildlife in question predicts less acceptance of lethal control has also been found cross-culturally for black bears (Liu & Sharp, 2018), wolves (Bruskotter et al., 2009), and jaguars and pumas (Engel et al., 2017). Together, this indicates that positive attitudes toward Australian wildlife would predict less support for lethal dingo control (see Figure 1, pathway *b*).

#### 1.9.1. Measuring Attitude towards Australian Wildlife

Attitude towards wildlife is typically measured using semantic-differential scales (e.g., bad vs. good, or harmful vs. beneficial; see Whitehouse□Tedd et al., 2021 for a review). However, a recent principal component analysis (Smith et al., submitted) classified attitudes toward Australian wildlife across five dimensions: Wildlife Ethics and Stewardship, Nature Affinity, Wildlife Apprehension, Wildlife Conservation, and Animal Welfare. Wildlife Ethics and Stewardship attitudes relate to how wildlife are valued and managed (e.g., for human benefit vs. the needs of wildlife). Nature Affinity attitudes encompass positive feelings, such as admiration, when connecting to wildlife and nature. Wildlife Apprehension attitudes reflect negative emotions towards wildlife, such as fear or dislike. Wildlife Conservation attitudes prioritise the interrelatedness of humans and non-humans, such that wildlife and ecosystems need to be proactively protected. Animal Welfare attitudes value the sentience and well-being of all animals. Smith et al. (submitted) found that Australian adults who agreed more strongly with the Wildlife Ethics and Stewardship (once reverse scored), Nature Affinity, Wildlife Conservation, and Animal Welfare subscales, and agreed less strongly with the Wildlife Apprehension subscale, demonstrated more positive attitudes overall toward Australian wildlife. While these dimensions have yet to be validated outside the scale development sample, their measure was used in the current study as it is the first attitudinal scale specific to Australian wildlife.

### 1.10. Pet Ownership

Pet ownership has also been associated with differences in support for the lethal management of wildlife. Australians report a high level (69%) of pet ownership (48% dogs; Animal Medicine Australia, 2022). Research has indicated that pet owners are generally pro-wildlife. In the United Kingdom, pet ownership (45% dogs) and positive attitudes towards pets predicted support for wildlife conservation, opposition to harmful wildlife management practices, and less support for prioritising human needs over that of animals (Shuttlewood et al., 2016). Similarly, when surveyed about a range of animals (e.g., chimpanzee, chicken, pig, dog), pet owners globally were less likely to endorse the use of animals for human benefit (e.g., research or pest control) than non-pet owners (Bradley et al., 2020). Together, this suggests that ownership or primary care of a dog as a pet may differentially predict support for lethal dingo control.

### 1.11. The Current Study

Following trends from other modern democratic nations, Australian animal-related organisations have been subject to social license pressure to ensure their practices align with public expectations (Hampton et al., 2020). The lethal control of dingoes has been publicly funded for over two centuries (Corbett, 2001). However, it was unknown whether contemporary Australians supported this practice. It was also unknown whether support for lethal dingo control was related to attitude towards Australian wildlife and social license pressure (as measured by public perceptions of procedural fairness, trust, and confidence). Findings from this study can inform the Australian government’s dingo and other wildlife policies and practices, and tailor attitudinal interventions and educational material for stakeholders, including the public. Furthermore, results can extend social license theory to the field of non-consumptive wildlife management. It can also add to human-animal intersection theory which includes recent movements advocating for non-human animal legal rights.

There were four aims of the current study. First, to assess Australian public support for the lethal control of dingoes. Second, to determine the predictive relationships between social license pressure, attitude towards Australian wildlife, and public support for lethal dingo control. Third, to determine if the relationship between social license pressure and public support for lethal dingo control was mediated by attitude towards Australian wildlife. Finally, to determine if there was a significant difference in support for lethal dingo control based on dog ownership. It was therefore hypothesised that:

Hypothesis 1 (H1): More Australians would oppose (rather than support) the lethal control of dingoes.

Hypothesis 2 (H2): Social license pressure would negatively predict Australian public support for the lethal control of dingoes (see Figure 1, pathway *c*).

Hypothesis 3 (H3): Social license pressure would positively predict attitude towards Australian wildlife (see Figure 1, pathway *a*).

Hypothesis 4 (H4): Attitude towards Australian wildlife would negatively predict public support for the lethal control of dingoes (see Figure 1, pathway *b*).

Hypothesis 5 (H5): The relationship between social license pressure and public support for the lethal control of dingoes would be mediated by attitude towards Australian wildlife (see Figure 1, pathway *ab*).

Hypothesis 6 (H6): Compared to individuals who have never owned a dog, past and/or current dog owners would be less supportive of lethal dingo control.

## 2. Material and methods

### 2.1. Participants

Eligible respondents were Australian residents aged 18 years and above. Of the 1102 responses, 112 (10%) were excluded as incomplete (104), duplicates (3), declined to consent (1) or failed the attention check (4). The 990 remaining respondents were aged between 18 and 83 years (*M_age_* = 50.0, *SD_age_* = 14.1), resided predominantly in Australia’s most populous states of Queensland, New South Wales, and Victoria (74%), and had spent most of their lives in urbanised areas (65%). Half the sample were women (52%) with tertiary qualifications (50%). Most respondents had experienced pet ownership (94% dog; 90% non-dog). More than two-thirds of respondents had interacted with live and captive dingoes in various settings, however only 39% knew that wild dog management in Australia included dingoes (van Eeden et al., 2021; survey item used with permission). The dog-owned and dog-never-owned sub-groups (*n* = 56 each) displayed similar demographic characteristics. See Appendix A for detailed demographic information.

### 2.2. Sampling Procedure

The current study was approved by Monash University Human Research Ethics Committee (Project Number: 24005). A voluntary Australian adult sample self-selected by responding to online advertisements on social media platforms (Facebook, LinkedIn, Twitter). A priori power analyses conducted to detect medium effect sizes as per Cohen’s (1988) conventions at power levels of 0.8 and alpha levels of .05 using G*Power (Version 3.1.9.7; Faul et al., 2009) and MedPower (Kenny, 2017). The analysed sample of 990 respondents was greater than the largest minimum requirement of 105 participants.

### 2.3. Measures

Measures were combined into one Qualtrics XM survey (Qualtrics LLC, 2020; see Appendix B). Participants were asked about their attitudes toward Australian wildlife, support for lethal dingo control, and social license pressure. They were also asked for their age, gender, state of residence, highest level of education, and type of area they had spent most of their lives in, dingo interactions, knowledge of the term “wild dogs”, and pet (dog and non-dog) ownership/primary care. All items were positively worded for face validity since prior social license scale development attempts found that negatively worded items confused respondents (Boutilier, 2017).

The novel items (support for lethal dingo control, social license pressure) were measured using 101-point (-50 to 50 including 0) continuous slider scales to provide maximum variability in the interval data (Chyung et al., 2018). Scale labels were only applied to mid (0) and endpoints (-50; 50) to minimise the risk of rounding bias that can occur when scores end in 0 or 5 (Maineri et al., 2021). The initial slider handle was set to 0 (neutral response) because endpoint or quartile positioning can result in satisficing bias (Bosch et al., 2019; Liu & Conrad, 2019; Toepoel & Funke, 2018).

#### 2.3.1. Support for Lethal Dingo Control

As research has shown that public awareness about dingo management is limited (van Eeden et al., 2021), participants were first advised that dingoes are legally managed in Australia, typically lethally (e.g., killing by poisoning). To elicit intuitive responses on the extent of opposition or support for lethal dingo control, the stimulus statement was presented around a neutral point of 0 (-50 = *strongly oppose*; 50 = *strongly support*). Distance from 0 indicated the strength of opposition or support. For analysis (see Figure 2), this was measured from 0 (*no support*) to 100 (*full support*), with the middle 20% of scores considered neutral in line with the midpoint option of 3 on a 5-point Likert scale (Chyung et al., 2017). Although there is no published reliability or validity data for this item, the use of oppose versus support question poles follows prior research into Australian public attitudes toward socially contentious topics (e.g., water distribution, Jackson et al., 2019; cigarette packet messaging, Brennan et al., 2021). Furthermore, single-item measures are appropriate when the construct is narrowly defined and unidimensional (Allen et al., 2022; Fuchs & Diamantopoulos, 2009). Our single-item measure was therefore suitable for the current study.

**Figure 2.**
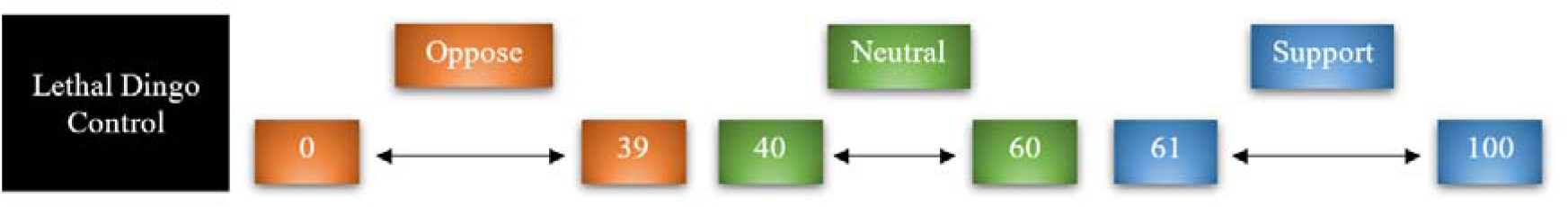
Support for Lethal Dingo Control: Scoring for Analysis Purposes.

#### 2.3.2. Social License Pressure

Social license pressure was measured by three 2-item subscales relating to procedural fairness, trust, and confidence of the Australian government’s management of dingoes. Example items included “The Australian government listens to and respects community opinions about dingo management” (Procedural Fairness), “The Australian government will be open and honest in relation to dingo management” (Trust), and “The Australian government does a good job managing dingoes” (Confidence). Trust Item 1, and both procedural fairness items, were adapted from social license models of mining in Australian, Chilean, and Chinese general population samples (Zhang et al., 2015; α = .74 - .96). Trust Item 2, and both confidence items, were adapted from principal components analyses of hunters’ acceptance, trust, and fairness perceptions of a waterfowl agency’s decisions in the United States (Schroeder et al., 2017; α = .88 - .96).

To elicit intuitive responses to the stimulus statements, and for consistency with the support for lethal dingo control item, social license pressure items were presented around a neutral point of 0 (-50 = *strongly disagree*; 50 = *strongly agree*). Distance from 0 indicated the strength of disagreement or agreement. For analysis (see Figure 3), items were measured from 0 (*strongly agree*) to 100 (*strongly disagree*), with the middle 20% of scores considered neutral (Chyung et al., 2017). Subscale items were summed for subscale totals (0 – 200). All items were summed for a composite social license pressure score (0 - 600). In the current study, internal consistency was excellent for the full scale (α = .93), and good for the subscales (α = .81 - .83).

**Figure 3.**
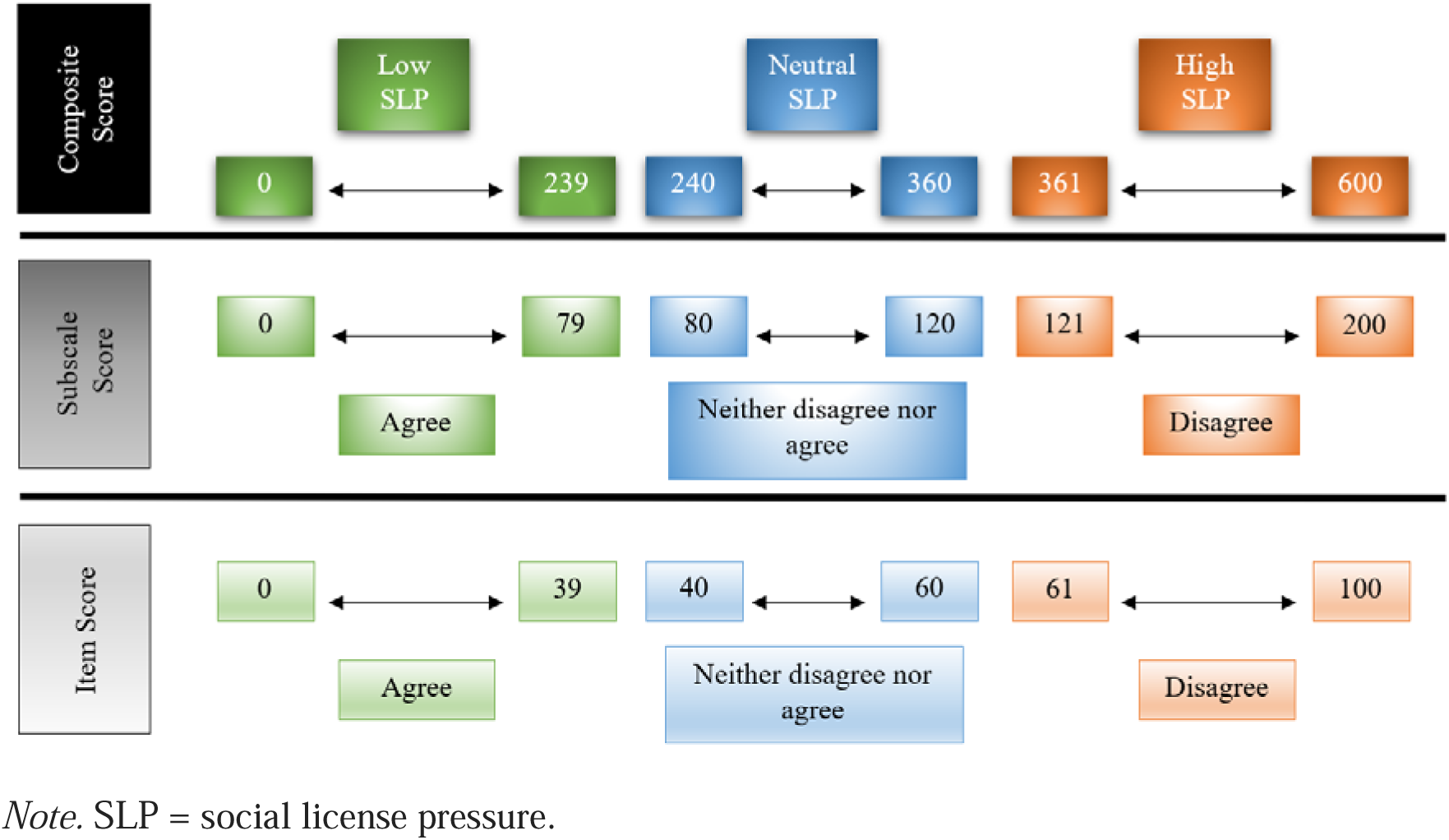
Social License Pressure: Scoring for Analysis Purposes.

#### 2.3.3. Attitude towards Australian Wildlife

Participant’s attitude towards Australian wildlife was measured by the Australian Attitudes to Wildlife Scale (AAWS; Smith et al., submitted; used with permission). The 30-item scale has five 6-item subscales rated on a 4-point Likert scale (1 = *strongly disagree*; 4 = *strongly agree*). Example items include “All forms of hunting are inhumane” (Wildlife Ethics and Stewardship), “I find wild animals fascinating” (Nature Affinity), “I don’t like animals” (Wildlife Apprehension), “Predators like dingoes keep nature in balance” (Wildlife Conservation), and “Animals have emotions” (Animal Welfare). After Wildlife Ethics and Stewardship Items 1, 2 and 3 were reverse scored, subscale items were summed for subscale totals (6 - 24). Higher subscale totals indicated positive attitudes, except for the Wildlife Apprehension subscale which indicated negative attitudes. A global attitude score (30 - 120) was calculated by subtracting the Wildlife Apprehension subscale from 30, then adding the other subscales (refer Smith et al, submitted). Higher global scores indicated more positive attitudes toward Australian wildlife.

Although newly developed, AAWS (Smith et al., submitted) was used because it is the first attitudinal scale specific to Australian wildlife. It demonstrated good internal consistency with the pilot adult Australian sample (*N* = 379, full scale α = .89, subscales α = .80 - .90). Construct validity was confirmed through correlations in the expected directions with participants’ self-reported behaviours, such as purchasing environmentally friendly products. In the current study, internal consistency was excellent for the full scale (α = .91), and acceptable for the subscales (α = .65 - .92).

### 2.4. Procedure

After viewing the survey advertisement, participants followed a hyperlink to the study’s explanatory statement. This detailed the study’s purpose, risks, benefits, eligibility criteria, and counselling support options. Participation was anonymous, confidential, voluntary, and available at any time/place with any internet-connected device. They could withdraw at any time before submission. Participants who consented and confirmed eligibility entered the survey. Those who did not consent or were ineligible were directed to the exit statement.

The survey (5 - 10 minutes to complete) was arranged in fixed blocks of AAWS (Smith et al., submitted), attention check (adapted from Prolific Academic Ltd, n.d.), support for lethal dingo control, social license pressure, and demographic items. This non-randomised block flow was adopted to enhance completer rates since it allowed a logical flow from broad (e.g., Australian wildlife) to specific (e.g., dingo control). Items within blocks (except demographic items) were randomised to minimise order effects. All items and blocks were force-choice responses. Demographic items included opt-out answers except for age because it was part of the eligibility criteria. The exit statement included a restatement of counselling support options.

### 2.5. Design and statistical analyses

This non-experimental study used cross-sectional survey data where the main variables were measured continuously. Although the AAWS (Smith et al., submitted) is an ordinal level Likert response scale, the current study treated it as interval data in line with current social science conventions (e.g., see review by Norman, 2010). A one-way chi-square test examined whether the observed frequency counts of oppose versus support scores (excluding neutral responses) on the lethal dingo control item were significantly different than expected (equal expected categories; H1). A mediation analysis tested the predictive relationships between social license pressure and support for lethal dingo control (H2; pathway *c*), between social license pressure and attitude towards Australian wildlife (H3; pathway *a*), and between attitude towards Australian wildlife and support for lethal dingo control (H4; pathway *b*). It also tested whether the relationship between social license pressure and support for lethal dingo control was mediated by attitude towards Australian wildlife (H5; pathway *ab*). Social license pressure was the predictor variable, measured by the composite score of procedural fairness, trust, and confidence items. Support for lethal dingo control was the outcome variable, measured by the score on the lethal dingo control item. Attitude towards Australian wildlife was the mediator variable, measured by the global score on the AAWS. An independent samples one-tailed *t*-test tested whether support for lethal dingo control differed by dog ownership (never owned vs. owned in past, currently or both; H6).

## 3. Results

Data cleaning was undertaken in Microsoft Excel (Office 365), and statistical analyses in IBM SPSS Statistics (Version 29). Details pertaining to data cleaning and assumption testing are available in Appendix C.

### 3.1. Preliminary Analyses

#### 3.1.1. One-Way Chi-Square Test (H1)

There were 520 participants (52.5%) whose score was categorised as opposing lethal dingo control (0 - 39), and 390 participants (39.4%) categorised as supporting (61 - 100). The 80 participants (8.1%) who gave neutral scores (40 - 60) were excluded from the analysis.

#### 3.1.2. Mediation Model (H2 – 5)

Given that variables demonstrated large standard deviations, significant skew, and bimodal distributions (see Appendix C), patterns across averaging methods (mean, median, mode) were considered to be a more accurate reflection of participants’ attitudes. For the subscale and composite levels of social license pressure, the increasing scores across averaging methods indicated that participants demonstrated attitudes of neutral to high social license pressure. Similarly, the decreasing score across averaging methods of support for lethal dingo control indicated that participants demonstrated neutral to no support for lethal dingo control. Attitude towards Australian wildlife global and subscales displayed smaller standard deviations, with averages demonstrating increasingly positive attitudes toward Australian wildlife, except for the Wildlife Ethics and Stewardship subscale. See Table 3 for further detail.

**Table 1.** Skewness, Kurtosis and Normality Test Results for Mediation Model.

| Parameter / Test | Social license<br>pressure | Attitudes to<br>Australian wildlife | Support for lethal<br>dingo control |
| --- | --- | --- | --- |
| Skewness |  |  |  |
| Statistic | -0.67 | 0.42 | 0.26 |
| $SE_{Skewness}$ | 0.08 | 0.08 | 0.08 |
| $Z_{Skewness}$ | -8.38 | 5.25 | 3.25 |
| Kurtosis |  |  |  |
| Statistic | -0.20 | -0.44 | -1.67 |
| $SE_{Kurtosis}$ | 0.16 | 0.16 | 0.16 |
| $Z_{Kurtosis}$ | -1.25 | -2.75 | -10.44 |
| Kolmogorov-Smirnov<br>statistic <sup>a</sup> | .12* | .08* | .21* |
| Shapiro-Wilks statistic | .92* | .97* | .80* |

*Note. N = 990.*
<sup>a</sup> Lilliefors Significance Correction.
\* $p < .001$ .

**Table 2.** Skewness, Kurtosis, and Normality Test Results for T-Test Analysis.

| Parameter / Test | Dog-Owned | Dog-Never-Owned |
| --- | --- | --- |
| Skewness |  |  |
| Statistic | 0.64 | -0.13 |
| $SE_{Skewness}$ | 0.32 | 0.32 |
| $Z_{Skewness}$ | 2.00 | -0.41 |
| Kurtosis |  |  |
| Statistic | -1.21 | -1.29 |
| $SE_{Kurtosis}$ | 0.63 | 0.63 |
| $Z_{Kurtosis}$ | -1.92 | -2.05 |
| Kolmogorov-Smirnov statistic ($p$)<sup>a</sup> | .28 (< .001) | .12 (.037) |
| Shapiro-Wilks statistic ($p$) | .78 (< .001) | .92 (.001) |

*Note.* $N = 112$ , $n = 56$ for each group.
<sup>a</sup>Lilliefors Significance Correction.

**Table 3.** Descriptive Statistics for Mediation Model.

| Variable | *M* | *SD* | Median | Mode | Min to Max |
| --- | --- | --- | --- | --- | --- |
| Social license pressure | 441.29 | 137.61 | 463.00 | 600 | 0 to 600 |
| Procedural fairness | 141.54 | 48.25 | 148.00 | 200 | 0 to 200 |
| Trust | 150.51 | 49.82 | 160.00 | 200 | 0 to 200 |
| Confidence | 149.24 | 50.33 | 159.00 | 200 | 0 to 200 |
| Attitudes to Australian wildlife | 97.75 | 13.16 | 99.00 | 106 | 49 to 120 |
| Ethics & stewardship <sup>a</sup> | 13.38 | 5.21 | 13.00 | 7 | 6 to 24 |
| Nature affinity <sup>a</sup> | 21.18 | 2.86 | 22.00 | 24 | 6 to 24 |
| Wildlife apprehension <sup>b</sup> | 8.22 | 2.08 | 8.00 | 6 | 6 to 17 |
| Wildlife conservation <sup>a</sup> | 20.63 | 3.33 | 22.00 | 24 | 6 to 24 |
| Animal welfare <sup>a</sup> | 20.78 | 3.18 | 22.00 | 24 | 6 to 24 |
| Support for lethal dingo control | 42.60 | 41.75 | 29.00 | 0 | 0 to 100 |

*Note.* $N = 990$ , Min = Minimum, Max = Maximum, Ethics & stewardship = Wildlife Ethics and Stewardship subscale.
<sup>a</sup> High score indicates more positive attitudes. <sup>b</sup> Low score indicates more positive attitudes.

All variables were significantly correlated, with medium to large effect size (*r* > .30, *p* < .001, two-tailed; Cohen, 1988). Social license pressure was positively correlated with attitudes to Australian wildlife (*r* = .34), and negatively correlated with support for lethal dingo control (*r* = - .41). This indicated that people who demonstrated attitudes consistent with high social license pressure were more likely to hold positive attitudes toward Australian wildlife, and less likely to support lethal dingo control. Attitudes to Australian wildlife negatively correlated with the strength of support for lethal dingo control (*r* = - .78). This revealed that individuals who supported lethal dingo control were less likely to hold positive attitudes toward Australian wildlife.

#### 3.1.3. Independent Samples T-Test (H6)

The participant group with dog-owning experience (dog-owned group) demonstrated attitudes consistent with high social license pressure across averaging methods, albeit with large standard deviations. While all three averages indicated opposition to lethal dingo control, the large standard deviation suggested that attitudes were varied. They held positive attitudes toward Australian wildlife at a global and subscale level, except for neutral attitudes regarding a utilitarian view of animals (i.e., Wildlife Ethics and Stewardship subscale).

The dog-never-owned group demonstrated weaker and more varied attitudes than the dog-owned group across all variables. They exhibited neutral or near-neutral attitudes of social license pressure, with composite and subscale scores at the boundary of neutral-high categories. Support for lethal dingo control was neutral based on mean and median score, however their mode score was the lowest possible no-support score. Both variables demonstrated large standard deviations, reflective of the varied attitudes among participants who had never owned a dog. Although they held positive attitudes towards Australian wildlife, their attitude was less strong than the dog-owned group. They also agreed with the utilitarian use of animals in the Wildlife Ethics and Stewardship subscale. See Table 4 for further detail.

**Table 4.** Descriptive Statistics for T-Test Analysis.

| Variable | *M* | *SD* | Median | Mode | Min to Max |
| --- | --- | --- | --- | --- | --- |
| Dog-Owned Group |  |  |  |  |  |
| Social license pressure | 462.77 | 121.57 | 468.00 | 600 | 165 to 600 |
| Procedural fairness | 145.13 | 47.75 | 146.00 | 200 | 0 to 200 |
| Trust | 158.07 | 40.23 | 162.00 | 200 | 35 to 200 |
| Confidence | 159.57 | 46.88 | 178.50 | 200 | 0 to 200 |
| Attitudes to Australian wildlife | 101.18 | 12.58 | 105.00 | 100 | 66 to 119 |
| Ethics & stewardship <sup>a</sup> | 14.48 | 5.00 | 15.00 | 14 | 6 to 24 |
| Nature affinity <sup>a</sup> | 21.77 | 2.52 | 23.00 | 24 | 13 to 24 |
| Wildlife apprehension <sup>b</sup> | 8.00 | 2.06 | 8.00 | 6 | 6 to 15 |
| Wildlife conservation <sup>a</sup> | 21.39 | 3.11 | 22.00 | 24 | 10 to 24 |
| Animal welfare <sup>a</sup> | 21.54 | 3.21 | 23.00 |  | 11 to 24 |
| Support for lethal dingo control | 33.36 | 38.74 | 10.00 | 0 | 0 to 100 |
| Dog-Never-Owned Group |  |  |  |  |  |
| Social license pressure | 370.21 | 115.51 | 356.50 |  | 143 to 575 |
| Procedural fairness | 117.88 | 42.78 | 107.00 |  | 43 to 200 |
| Trust | 131.09 | 46.03 | 132.50 | 200 | 30 to 200 |
| Confidence | 121.25 | 40.41 | 121.00 |  | 48 to 200 |
| Attitudes to Australian wildlife | 93.77 | 11.45 | 95.00 | 95 | 59 to 114 |
| Ethics & stewardship <sup>a</sup> | 12.91 | 3.88 | 13.50 | 14 | 6 to 22 |
| Nature affinity <sup>a</sup> | 20.09 | 2.96 | 20.00 | 24 | 11 to 24 |
| Wildlife apprehension <sup>b</sup> | 9.75 | 2.76 | 9.00 | 8 | 6 to 17 |
| Wildlife conservation <sup>a</sup> | 20.23 | 2.82 | 21.00 |  | 11 to 24 |
| Animal welfare <sup>a</sup> | 20.29 | 2.74 | 21.00 |  | 12 to 24 |
| Support for lethal dingo control | 49.30 | 33.66 | 52.50 | 0 | 0 to 100 |

Note. N = 112 (*n* =56 per group), Min = Minimum, Max = Maximum, Ethics & stewardship = Wildlife Ethics and Stewardship subscale. Blank cells indicate that there were multiple modes.
<sup>a</sup> High score indicates more positive attitudes. <sup>b</sup> Low score indicates more positive attitudes.

### 3.2. Inferential Statistics

#### 3.2.1. One-Way Chi-Square Test

A one-way chi-square test assessed whether the observed frequency counts of those who opposed versus supported lethal dingo control were significantly different from equal expected categories (H1). The chi-square test was significant, χ^2^ (1, *N* = 910) = 18.57, *p* < .001, with a small effect size (ω = 0.14; Cohen, 1988). With positive residuals (observed frequencies minus expected frequencies) for the opposing category, and negative residuals for the supporting category, results indicated that more Australians opposed (rather than supported) the lethal control of dingoes.

#### 3.2.2. Mediation Model

A mediation model (PROCESS macro version 4.2, Model 4; Hayes, 2022), with 5,000 bootstrapped samples and an H3-heteroscedastic-consistent approach, examined the predictive relationships between social license pressure, attitude towards Australian wildlife, and support for lethal dingo control (H2; H3; H4), and whether attitude towards Australian wildlife mediated the relationship between social license pressure and support for lethal dingo control (H5). All predictive pathways were significant (*p* < .001). All effect sizes were assessed according to Cohen’s (1988) conventions. Social license pressure significantly accounted for variance in attitude towards Australian wildlife (pathway *a*), *F*(1, 988) = 106.61 with a small effect size (*f^2^* = 0.13). Social license pressure also significantly accounted for variance in support for lethal dingo control (pathway *c*), *F*(1, 988) = 179.26, with a medium effect size (*f^2^* = 0.20). There was support for mediation with a significant indirect effect (pathway *ab*) of social license pressure predicting support for lethal dingo control through attitude towards Australian wildlife, *F*(2, 987) = 1170.66, with a large effect size (*f^2^* = 1.73). See Table 5 for further detail.

**Table 5.** Mediation Model Results.

| Effect | $B$ | 95% CI for $B$ | | SE $B$ | $\beta$ | $p$ | $R^2$ |
| --- | --- | --- | --- | --- | --- | --- | --- |
| | | $LL$ | $UL$ | | | | |
| Total effect (pathway $c$) | -0.12 | -0.14 | -0.11 | 0.01 | -.41 | .000 | .17 |
| Pathway $a$ | 0.03 | 0.03 | 0.04 | 0.00 | .34 | .000 | .12 |
| Pathway $b$ | -2.31 | -2.43 | -2.19 | 0.06 | -.73 | .000 | |
| Direct effect (pathway $c'$) | -0.05 | -0.06 | -0.04 | 0.01 | -.16 | .000 | |
| Indirect effect (pathway $ab$) | -0.08 | -0.09<sup>a</sup> | -0.06<sup>a</sup> | 0.01<sup>b</sup> | -.25<sup>c</sup> | | .64 |

*Note.* $N = 990$ . CI = confidence interval; $LL$ = lower limit; $UL$ = upper limit.
<sup>a</sup> Bootstrapped confidence interval (BCI). <sup>b</sup> Bootstrapped standard error (BSE).
<sup>c</sup> 95% BCI [-0.29, -0.21], BSE ( $\beta$ ) = 0.02.

#### 3.2.3. Independent Samples T-Test (H6)

An independent samples t-test with group sizes equalised at 56 was undertaken to compare support for lethal dingo control between those who had never owned a dog, and those who currently and/or previously owned a dog. Results indicated that the dog-never-owned group (*M* = 49.30, *SD* = 33.66) gave higher (i.e., more favourable) ratings of support for lethal dingo control than those in the dog-owned group (*M* = 33.36, *SD* = 38.74), *t*(110) = 2.33, *p* = .011, one-tailed, equal variances assumed. The mean difference of 15.95 units of support for lethal dingo control was significant, 95% CI [2.36, 29.54], with small effect size (*d* = 0.44; Cohen, 1988).

## 4. Discussion

The lethal control of dingoes has been publicly funded since Australia was colonised with few examinations of whether this aligns with contemporary Australian public opinion. The current study explored the extent of support for lethal dingo control in a general sample of Australian adults. Further, we explored how support was related to social license pressure, attitudes towards Australian wildlife, and dog ownership. The results supported all hypotheses. More of our sample opposed (rather than supported) lethal dingo control. Social license pressure negatively predicted support for lethal dingo control, and positively predicted attitude towards Australian wildlife. Attitude towards Australian wildlife negatively predicted support for lethal dingo control. The relationship between social license pressure and support for lethal dingo control was mediated by attitude towards Australian wildlife. Dog owners (past and/or current) were less supportive of lethal dingo control than individuals who had never owned a dog.

### 4.1. Australian Public Support for Lethal Dingo Control

Our result that more Australians opposed lethal dingo control provides novel evidence of how the Australian public views dingo management. It extends van Eeden et al.’s (2021) findings that Australians held positive attitudes about dingoes and disapproved of all lethal methods to control dingoes. Given that 65% of our respondents had spent most of their lives in urbanised areas (where dingoes do not reside), our result also aligns with evidence that individuals with greater geographical distance from wolf populations demonstrated more positive attitudes towards the species and their conservation (Karlsson & Sjöström, 2007; Williams et al., 2002). Our result therefore adds evidence about dingoes to the existing human-wildlife and human-canid interaction literature.

While the exploratory nature of the current research precludes any direct inferences as to why there is public opposition to lethal dingo control, our finding is consistent with broader changes in Australian attitudes about animals. Research has found that contemporary Australians recognise the sentience of animals (Webster, 2022), prioritise animal welfare outcomes (Chen, 2016), and disapprove of animal and wildlife use for human benefit (Jacobs et al., 2022). Such attitudes and values have been strong enough to result in the application of social license pressure (e.g., protest behaviours) against previously accepted animal-use industries (e.g., greyhound racing, see Markwell et al., 2017). The desire to prevent lethal harm to dingoes may be another way for Australians to express their pro-animal values. Further research (e.g., qualitative research) is encouraged to explore the underlying reasons for public opposition to lethal dingo control.

### 4.2. Social License Pressure, Attitude towards Australian Wildlife, and Public Support for Lethal Dingo Control

The current study extends and links understanding within the human-animal interaction, social license, and dingo evidence base. For instance, our finding that individuals who demonstrated attitudes of higher social license pressure were less likely to support lethal control is novel for dingoes. This is consistent with previous findings that non-consumptive wildlife culling ceased or was threatened when public disapproval was high, including for koalas (Drijfhout et al., 2020) and brumbies (Nimmo et al., 2007). Additionally, our finding that individuals with more positive attitudes toward Australian wildlife were less likely to support lethal dingo control aligns with cross-cultural evidence about national or regional species, such as black bears in the United States (Liu & Sharp, 2018), and jaguars and pumas in Brazil (Engel et al., 2017). This also aligns with evidence that compared to individuals from six other nationalities, Australians demonstrated the most egalitarian attitudes toward wildlife, and most strongly disapproved of wildlife hunting (Jacobs et al., 2022). Our results, therefore, contribute to the integration of evidence about attitudes toward wildlife, social license, and dingoes.

These results also add new evidence to our understanding of attitudes toward wildlife and their management. First, individuals who displayed attitudes of stronger social license pressure were more likely to have positive attitudes toward Australian wildlife. Second, that attitude towards Australian wildlife mediated the relationship between social license pressure and support for lethal dingo control. While these findings are novel, they are not unexpected. This is because many Australians distrust their government, with 44% reporting no or low trust in the national government (Organisation for Economic Cooperation and Development [OECD], 2022). Australians also value the welfare of animals (Chen, 2016) and believe in the equal treatment of humans and wildlife (Jacobs et al., 2022). This relates to their distrust of animal-use industries, which often demonstrate secrecy and a lack of accountability around animal welfare outcomes (e.g., see Hampton et al., 2020). Our study provides novel evidence that social license pressure and attitudes towards wildlife play important roles in explaining support or opposition to lethal control in wildlife management.

The current study provides evidence that contemporary Australians generally demonstrate pro-wildlife conservation values and consider lethal management practices to be inappropriate. However, there was considerable heterogeneity in attitudes about lethal dingo control. This may be related to the unequal distribution of wildlife management costs and benefits. For example, because rural communities and agricultural industries often bear the costs of coexisting with wildlife, they demonstrate less support for wildlife conservation compared with urban populations (Jordan et al., 2020). While further research is required to explore the underpinning beliefs and mechanisms of our results, multiple human factors must be considered for decisions about wildlife management to be publicly supported.

### 4.3. Dog Ownership

Our finding that dog owners were less supportive of lethal dingo control than those who had never experienced dog ownership is consistent with previous studies. Pet ownership has previously predicted opposition to management practices that harm wildlife (Shuttlewood et al., 2016), and less support for using animals in research or for pest control (Bradley et al., 2020). While our study design does not support direct inferences as to why dog ownership changes support for lethal wildlife control, research has indicated that pet owners assign stronger affective attitudes (e.g., love) and weaker utilitarian value (e.g., animal-use for human benefit) toward animals (Serpell, 2004). This suggests that affection for and familiarity with a domestic animal species for companionship (e.g., dogs as pets) may be related to disapproval of harm for their wild relation (e.g., dingoes). Similarly, in line with animal moralistic attitudes (Kellert, 1984), pet owners may ascribe a higher intrinsic value to animals, and view animal harm as morally wrong.

### 4.4. Implications of the Current Study

#### 4.4.1. Theoretical

The current study provides support for the applicability of social license theory to wildlife management. Such application is currently nascent and inconsistent (e.g., see Garnett et al., 2018; Kendal & Ford, 2018). Our results provide a novel link between social license and support for lethal control in the context of dingoes. They also offer a more refined understanding of that relationship through the inclusion of attitude towards Australian wildlife as a mediator variable. In addition, this gives preliminary evidence of three facets relevant to measuring social license pressure in a wildlife management context. Together, our results can support the development of validated social license models for wildlife management and other animal-based interactions, where none currently exist. This has implications for policy.

#### 4.4.2. Policy

Validated social license tools for wildlife management can assist with predicting a potential loss of social license, and tracking changes in social license over time, so policy can be developed proactively. Once the public becomes aware (often through media activism) of an animal-use practice that they oppose, the loss of social license can quickly force industry change (Hampton et al., 2020). For example, when California banned the importation of kangaroo products due to animal welfare concerns, there were immediate deleterious effects on the $200 million annual retail value of kangaroo products (March, 2015); kangaroo skins became nearly worthless, and industry employment declined (Wilson & Edwards, 2019). Although dingo and kangaroo culling are not directly comparable because kangaroo culling has commercial (e.g., sale of meat and skins) and non-commercial purposes (e.g., reducing competition for livestock feed), a loss of social license for lethal dingo control would nonetheless have a direct impact. For example, the employment of dingo controllers, and the cashflow of businesses that manufacture dingo bait. It is therefore important that validated social license instruments for wildlife management be developed to predict challenges and changes to social license, so policies can be implemented to minimise the risks to Australian industries and employment.

Similarly, the current study may inform education and dialogue with individual stakeholders (e.g., agricultural producers, policy makers) who are unaware or resistant to acknowledging the deleterious effects on industry from a loss of social license. Our results contribute to the small but growing evidence base that social license for lethal dingo control is tenuous in Australia. If public opposition to lethal dingo control were mobilised through media activism, this would fundamentally challenge livestock grazing that covers more than half the continent (ABARES, 2016). The loss of social license has already changed lethal management practices in Australia for brumbies (Nimmo et al., 2007) and kangaroos (Hampton & Teh-White, 2019). In the case of feral camel control, attention to social license principles has yielded diverse positive outcomes, such as biodiversity protection and grazing soil improvement, while still reducing feral camel densities (Hart & Bubb, 2016). Acknowledging the need for social license and public support is therefore crucial for sustainable and successful wildlife management policies.

Importantly, alternative strategies for livestock protection and non-lethal methods for dingo management exist but have received little uptake or further research investment (e.g., Smith et al., 2020; Smith et al., 2021; van Bommel and Johnson, 2023). Given that social license for lethal control of dingoes is currently tenuous, further exploration of alternative methods is necessary (Fancourt et al., 2021). Because societal attitudes now value animal welfare and seek its assurance, governments are likely to gain or lose support from the public on human-animal interaction issues such as this in the future (Yin et al., 2023). Further research to better elucidate the role of intuitive judgmental heuristics and individual value orientations when stakeholders form impressions about the distinct framing of livestock protection and wildlife control practices would be valuable (Goto Gray et al., 2020). Social license assessment tools such as those used in the current study can indicate where significant change in both human-animal interaction policy and practice is expected by modern Australians.

#### 4.4.3. Informing attitudinal interventions and mental health support programs

Traditional deficit model knowledge-transfer interventions often fail to effectively change behaviour relating to animal management (see McLeod et al., 2015). Our mediation results may assist with the design of attitudinal interventions. When mediator variables are changeable with a clear directional relationship to the outcome variable, they may be targeted to elicit change in the outcome variable (MacKinnon et al., 2007). Weaker attitudes are easier to change than stronger ones, and when the topic is personally important, it motivates attitude-expressive behaviour, such as contacting government representatives (Howe & Krosnick, 2017). Together with the finding that attitude towards Australian wildlife mediates the relationship between social license pressure and support for lethal dingo control, this suggests that an attitudinal intervention would be effective. For example, advocates for nonlethal dingo management may target individuals (e.g., via a public education campaign) who indicate that wildlife management is important to them, and currently express neutral or slightly negative attitudes toward Australian wildlife. Our results suggest that if these attitudes toward Australian wildlife became more positive, then these individuals may also shift to opposing (rather than supporting) lethal dingo control.

Our results could also assist in the treatment of distressed individuals engaged in sustainability and animal-care work or volunteerism. These individuals include researchers (King & Zohny, 2022), scientists (Conroy, 2019), veterinary staff (White et al., 2021), veterinary students (Littlewood et al., 2020), laboratory animal personnel (LaFollette et al., 2020), and wildlife carers (Yeung et al., 2017). Their strong commitment to the care of animals and the environment means they are especially vulnerable to empathy-based distress (e.g., compassion fatigue, secondary traumatic stress, vicarious traumatisation; see review by Rauvola et al., 2019), or ecological anxiety and grief (see review by Pihkala, 2024). Our finding that most Australians demonstrate attitudes consistent with strong social license pressure and low support of lethal dingo control may provide a form of distal social support. This is important because a sense of shared identity and community can facilitate empowerment and meaning-making for those involved in collective action (Clayton, 2018; Drury & Reicher, 2009). For example, climate scientists report feeling supported by contact with concerned members of the public (Richardson, 2015) despite being consistently overwhelmed by the scope of the problems (Conroy, 2019; Holcombe et al., 2016). While the development of “ecotherapy” is in its infancy (Cunsolo & Ellis, 2018), the social support implicit in our results may be beneficial in the development of resilience strategies for individuals engaged in animal-care and sustainability work.

### 4.4. Strengths and Limitations of the Current Study

A strength of the current study was its novel and much-needed focus on the Australian general public in assessing support for lethal dingo control. Existing discussions regarding dingo control in Australia typically focus on the economic and social cost to agricultural producers from dingo predation of livestock (see Australian Wool Innovation Ltd, 2020). While the costs to agricultural communities are real and extend beyond a monetary value (Ecker et al., 2017), such a focus excludes other stakeholders (e.g., general public), and other perspectives (e.g., sustainable wildlife use for non-monetary matters, such as ecological and cultural significance). Given that all of the Australian public funds dingo control through their taxes, and elects government representatives that vote on wildlife management policies, their voice should be equally included. Exclusion of societal opinions has demonstrably been to the detriment of various animal and wildlife-reliant industries, which have faced increasing social license pressure (e.g., see Hampton & Teh-White, 2019). Our study’s recognition of the importance of the attitudes, values, and expectations of the Australian public is crucial to understanding how wildlife management (and more broadly, human-animal interactions) can be sustainable and successful.

Another strength of the current study was our innovative adaption of validated tools from the broader social license literature, which has only recently been applied to wildlife management (Kendal & Ford, 2018). Our measurement of social license pressure through the facets of procedural fairness, trust, and confidence with good internal consistency provides preliminary evidence of their importance to understanding support for wildlife management. Additionally, our delineation between trust and confidence avoids the conflation seen in other wildlife attitudinal studies (e.g., Engel et al., 2016). Our measurement of social license pressure complements and extends other findings that trust between stakeholder groups, perceptions of effectiveness, fair representation, and pluralistic consultation are crucial to the successful management of large carnivores (Sjölander-Lindqvist et al., 2015), and large mammals (Hart & Bubb, 2016). Our study, therefore, demonstrates one way of extending social license theory to wildlife management.

A final strength of the current study was its good response rate, sufficient statistical power for all analyses, and adequate sampling representativeness compared to census data. Our sample size was large enough for all analyses to be sufficiently powered to detect significant effects. The strong response rate and limited drop-out rate suggest that our survey was novel enough to attract respondents yet parsimonious enough to avoid survey fatigue. Our convenience sample was also comparable to the latest Australian census data. Our (binary) gender split was reasonably even (45% men; 51% women). This is comparable to the 2021 Australian census (49% men; 51% women; ABS, 2022b), and does not demonstrate the gender effect often seen in social research (Becker, 2022). Additionally, census data (ABS, 2022b) indicated that 78% of Australians lived on the Eastern seaboard, which is comparable to our sample (75%). Furthermore, 28% of Australians are classified as residing in rural and remote areas (ABS, 2022a). Although we did not capture the type of area of current residence, participants did self-classify the type of area they had spent most of their lives in. With 34% of our sample having spent most of their lives in rural, remote, or very remote areas, this suggests that our sample had a higher level of lived experience in non-urban areas comparable to the broader Australian population. Such indicators of sampling validity enable greater confidence in the generalisability of our results.

Whilst our adaption of social license pressure from the broader social license literature was innovative, our measure has not yet been further validated. Although the wording of items was specific to the government’s management of dingoes, we did not include additional items to differentiate perceptions of procedural fairness, trust, and confidence in the government in general. Research indicates that perceptions of government may differ from general to specific contexts. For example, although 44% of Australians reported distrusting the national Australian government (OECD, 2022), 72% of Australians agreed that because of the federal government’s handling of the COVID-19 pandemic, they now considered the government to be generally trustworthy (Goldfinch et al., 2021). Therefore, it is unknown if our social license pressure measure inadvertently captured generalised anti-government sentiment rather than specific perceptions relating to the government’s management of dingoes.

Such a context difference (general vs. specific) in perceptions of government must be interpreted with caution. This is because there are limited parallels between the perception of government policies that directly affect a small number of people daily (e.g., wildlife management), compared to those that affect all members of a population (e.g., a public health crisis that restricts daily activities). Further, the “rallying around the flag” effect for government and public institutions seen in times of crises (Baekgaard et al., 2019; Schuman, 1975) is unlikely to translate to longstanding wildlife management policies that do not directly impact the majority of Australia’s urbanised population. Further research, such as factor analyses, would be required to confirm the content and construct validity of our measure of social license pressure. Nonetheless, our pilot measure with good internal consistency may be considered a sound point from which future research may explore social license pressure in relation to wildlife management and other human-animal interactions.

We are unable to infer if our results are sufficiently representative of those who typically have direct contact with dingoes (e.g., rural communities, agricultural producers). Although attempts were made to share the study invitation widely across Australia, including deliberate sharing with online agricultural community and rural-focussed groups, we have no data as to the relative urban-rural uptake. To reduce the risk of survey fatigue, we did not ask participants if they identified with any stakeholder groups, such as hunters or agricultural producers. Agricultural or hunter stakeholder identification and those with a rural upbringing have been associated with more negative attitudes toward coexistence with predator wildlife compared to urban populations (Landon et al., 2019; Vaske et al., 2022). While our sample had a higher level of rural lived experience (34%) than those in comparable Australian studies (e.g., 19%; van Eeden et al., 2021), and census data (e.g., rural and remote residence 28%; ABS, 2022a), it is unknown if our sample was sufficiently representative of individuals who have contact with dingoes as part of their daily lives. To ensure more targeted sampling, future research should seek the support of rural organisations (e.g., National Farmer’s Federation) to distribute study invitations, and ask participants about any specific stakeholder identifications.

### 4.5. Future Research

In addition to the aforementioned suggestions for future research, researchers are encouraged to explore other variables that may influence support or opposition to lethal dingo control. These may include knowledge, familiarity, length of pet ownership time, and closeness of pet breeds to dingoes. Research has shown that higher factual knowledge about wolves correlates with more positive attitudes toward them (Arbieu et al., 2019). However, it is unknown whether this effect exists for dingoes, and how accurate the knowledge of the Australian public is about dingoes. Similarly, the degree of familiarity with wolves was associated with differing levels of support for wolf conservation (Karlsson & Sjöström, 2007). This suggests that there is utility in examining how interacting with dingoes in real life versus media sources, or with wild dingoes versus captive dingoes, changes support for coexistence with dingoes. Following our result of differences in support for lethal dingo control based on dog ownership, further research could explore the possible moderating effects of length of dog ownership time, and how close in physical appearance a breed of pet dog is to a dingo. Likewise, exploring whether differences in support for lethal dingo control are demonstrated if the pet is not a dog. Further nuanced research can assist with the facilitation of more sustainable and less conflictual human-wildlife outcomes.

## 5. Conclusion

Dingo-livestock conflict in Australia remains unabated and unresolved after two centuries of publicly funded extermination efforts. The current study provided a much-needed assessment of attitudes toward dingoes, Australian wildlife, and social license for lethal control in a large convenience sample of Australian adults that compared well to census data. Our results indicated that support for lethal dingo control was heterogeneous (although low on average), predicted by social license pressure and attitude towards Australian wildlife, and differed by dog-ownership status. We also demonstrated that attitude towards Australian wildlife explained a large proportion of variance in the relationship between social license pressure and support for lethal dingo control. Additionally, we provided preliminary evidence of the importance of procedural fairness, trust, and confidence in measuring social license pressure for the wildlife management field. With human and dingo population trends showing no sign of easing, human-wildlife conflict will only increase unless new approaches are taken. These approaches should be transdisciplinary and incorporate evidence from relevant social and applied psychological theories, such as social license theory. They should also be empirically supported, including only wildlife management strategies shown to be both humane for wildlife and effective for successful coexistence. We recommend that future research involves all stakeholders, including the general public, in order to improve outcomes for humans and wildlife on this ever-evolving planet.

## Author contributions (CRediT taxonomy)

AIP: Conceptualization (Equal); Methodology (Lead); Data curation; Formal analysis; Investigation; Writing - original draft (Lead); Writing - review & editing.

MLC: Conceptualization (Equal); Methodology; Project administration; Supervision; Writing - review & editing.

## Acknowledgements

An earlier version of this work was submitted as a thesis in partial fulfilment for the Graduate Diploma in Psychology Advanced at Monash University, informed by feedback from Dr. Ryan Anderson, Dr. Rhys Luckey, and Dr. Pamela Pensini. We are grateful to Dr. Bradley Smith from Central Queensland University for the use of the Attitudes to Australian Wildlife Scale and the feedback from the journal’s anonymous reviewers.

## Declaration of interests

⍰ The authors declare that they have no known competing financial interests or personal relationships that could have appeared to influence the work reported in this paper.

⍰ The authors declare the following financial interests/personal relationships which may be considered as potential competing interests:

## Funding

The authors received no funding for this study.

## Ethical oversight

The Monash University Human Research Ethics Committee approved this study (MUHREC Approved Project: 24005).

## Appendix A Demographic Characteristics of Analysed Sample

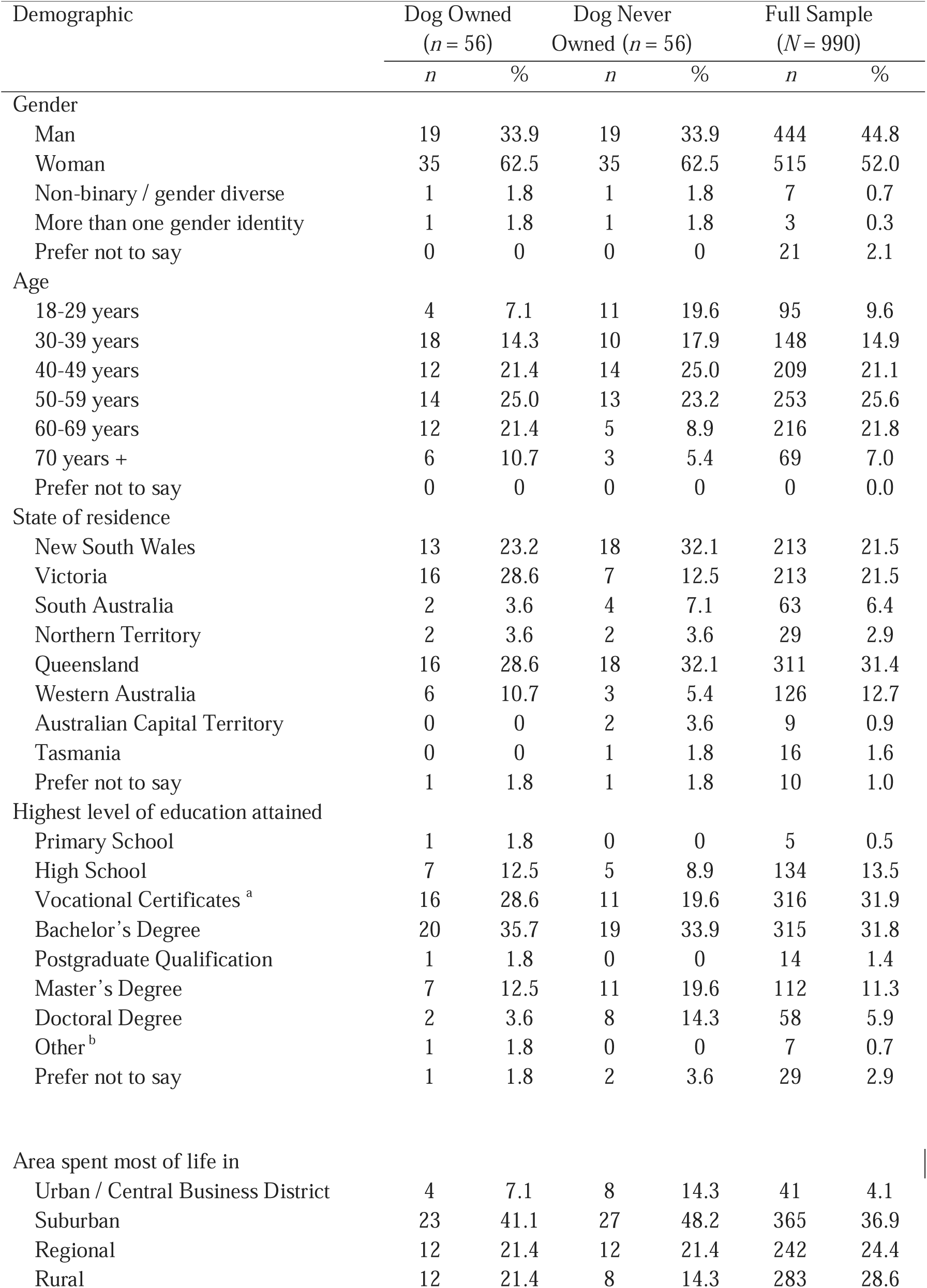
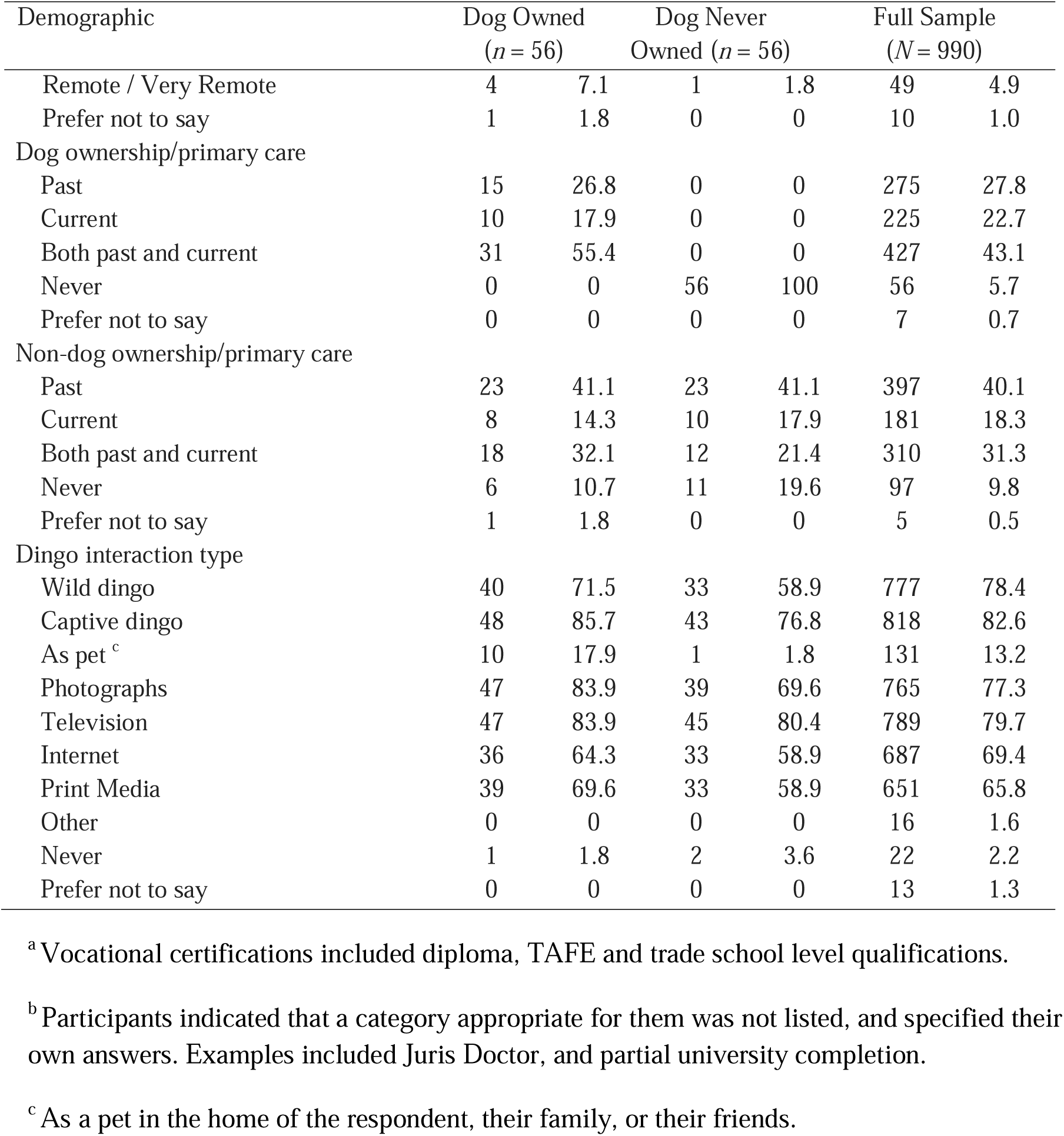

Demographic Characteristics of Analysed Sample
| Demographic | Dog Owned<br>(*n* = 56) |  | Dog Never<br>Owned (*n* = 56) |  | Full Sample<br>(*N* = 990) |  |
| --- | --- | --- | --- | --- | --- | --- |
|  | *n* | % | *n* | % | *n* | % |
| Gender |  |  |  |  |  |  |
| Man | 19 | 33.9 | 19 | 33.9 | 444 | 44.8 |
| Woman | 35 | 62.5 | 35 | 62.5 | 515 | 52.0 |
| Non-binary / gender diverse | 1 | 1.8 | 1 | 1.8 | 7 | 0.7 |
| More than one gender identity | 1 | 1.8 | 1 | 1.8 | 3 | 0.3 |
| Prefer not to say | 0 | 0 | 0 | 0 | 21 | 2.1 |
| Age |  |  |  |  |  |  |
| 18-29 years | 4 | 7.1 | 11 | 19.6 | 95 | 9.6 |
| 30-39 years | 18 | 14.3 | 10 | 17.9 | 148 | 14.9 |
| 40-49 years | 12 | 21.4 | 14 | 25.0 | 209 | 21.1 |
| 50-59 years | 14 | 25.0 | 13 | 23.2 | 253 | 25.6 |
| 60-69 years | 12 | 21.4 | 5 | 8.9 | 216 | 21.8 |
| 70 years + | 6 | 10.7 | 3 | 5.4 | 69 | 7.0 |
| Prefer not to say | 0 | 0 | 0 | 0 | 0 | 0.0 |
| State of residence |  |  |  |  |  |  |
| New South Wales | 13 | 23.2 | 18 | 32.1 | 213 | 21.5 |
| Victoria | 16 | 28.6 | 7 | 12.5 | 213 | 21.5 |
| South Australia | 2 | 3.6 | 4 | 7.1 | 63 | 6.4 |
| Northern Territory | 2 | 3.6 | 2 | 3.6 | 29 | 2.9 |
| Queensland | 16 | 28.6 | 18 | 32.1 | 311 | 31.4 |
| Western Australia | 6 | 10.7 | 3 | 5.4 | 126 | 12.7 |
| Australian Capital Territory | 0 | 0 | 2 | 3.6 | 9 | 0.9 |
| Tasmania | 0 | 0 | 1 | 1.8 | 16 | 1.6 |
| Prefer not to say | 1 | 1.8 | 1 | 1.8 | 10 | 1.0 |
| Highest level of education attained |  |  |  |  |  |  |
| Primary School | 1 | 1.8 | 0 | 0 | 5 | 0.5 |
| High School | 7 | 12.5 | 5 | 8.9 | 134 | 13.5 |
| Vocational Certificates <sup>a</sup> | 16 | 28.6 | 11 | 19.6 | 316 | 31.9 |
| Bachelor's Degree | 20 | 35.7 | 19 | 33.9 | 315 | 31.8 |
| Postgraduate Qualification | 1 | 1.8 | 0 | 0 | 14 | 1.4 |
| Master's Degree | 7 | 12.5 | 11 | 19.6 | 112 | 11.3 |
| Doctoral Degree | 2 | 3.6 | 8 | 14.3 | 58 | 5.9 |
| Other <sup>b</sup> | 1 | 1.8 | 0 | 0 | 7 | 0.7 |
| Prefer not to say | 1 | 1.8 | 2 | 3.6 | 29 | 2.9 |
| Area spent most of life in |  |  |  |  |  |  |
| Urban / Central Business District | 4 | 7.1 | 8 | 14.3 | 41 | 4.1 |
| Suburban | 23 | 41.1 | 27 | 48.2 | 365 | 36.9 |
| Regional | 12 | 21.4 | 12 | 21.4 | 242 | 24.4 |
| Rural | 12 | 21.4 | 8 | 14.3 | 283 | 28.6 |
| Remote / Very Remote | 4 | 7.1 | 1 | 1.8 | 49 | 4.9 |
| Prefer not to say | 1 | 1.8 | 0 | 0 | 10 | 1.0 |
| Dog ownership/primary care |  |  |  |  |  |  |
| Past | 15 | 26.8 | 0 | 0 | 275 | 27.8 |
| Current | 10 | 17.9 | 0 | 0 | 225 | 22.7 |
| Both past and current | 31 | 55.4 | 0 | 0 | 427 | 43.1 |
| Never | 0 | 0 | 56 | 100 | 56 | 5.7 |
| Prefer not to say | 0 | 0 | 0 | 0 | 7 | 0.7 |
| Non-dog ownership/primary care |  |  |  |  |  |  |
| Past | 23 | 41.1 | 23 | 41.1 | 397 | 40.1 |
| Current | 8 | 14.3 | 10 | 17.9 | 181 | 18.3 |
| Both past and current | 18 | 32.1 | 12 | 21.4 | 310 | 31.3 |
| Never | 6 | 10.7 | 11 | 19.6 | 97 | 9.8 |
| Prefer not to say | 1 | 1.8 | 0 | 0 | 5 | 0.5 |
| Dingo interaction type |  |  |  |  |  |  |
| Wild dingo | 40 | 71.5 | 33 | 58.9 | 777 | 78.4 |
| Captive dingo | 48 | 85.7 | 43 | 76.8 | 818 | 82.6 |
| As pet <sup>c</sup> | 10 | 17.9 | 1 | 1.8 | 131 | 13.2 |
| Photographs | 47 | 83.9 | 39 | 69.6 | 765 | 77.3 |
| Television | 47 | 83.9 | 45 | 80.4 | 789 | 79.7 |
| Internet | 36 | 64.3 | 33 | 58.9 | 687 | 69.4 |
| Print Media | 39 | 69.6 | 33 | 58.9 | 651 | 65.8 |
| Other | 0 | 0 | 0 | 0 | 16 | 1.6 |
| Never | 1 | 1.8 | 2 | 3.6 | 22 | 2.2 |
| Prefer not to say | 0 | 0 | 0 | 0 | 13 | 1.3 |

<sup>a</sup>Vocational certifications included diploma, TAFE and trade school level qualifications.
<sup>b</sup>Participants indicated that a category appropriate for them was not listed, and specified their own answers. Examples included Juris Doctor, and partial university completion.
<sup>c</sup> As a pet in the home of the respondent, their family, or their friends.

## Appendix B Study Survey

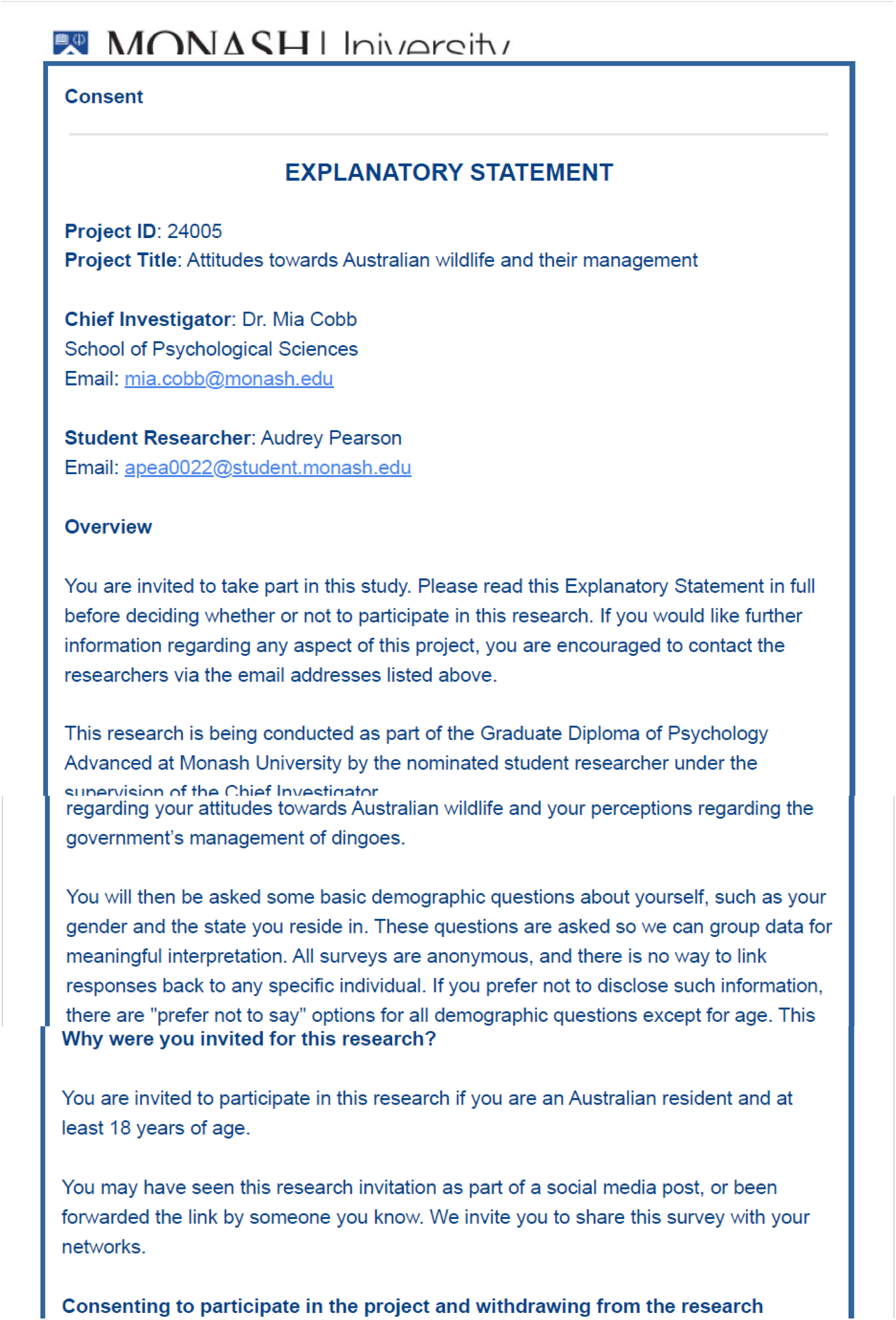
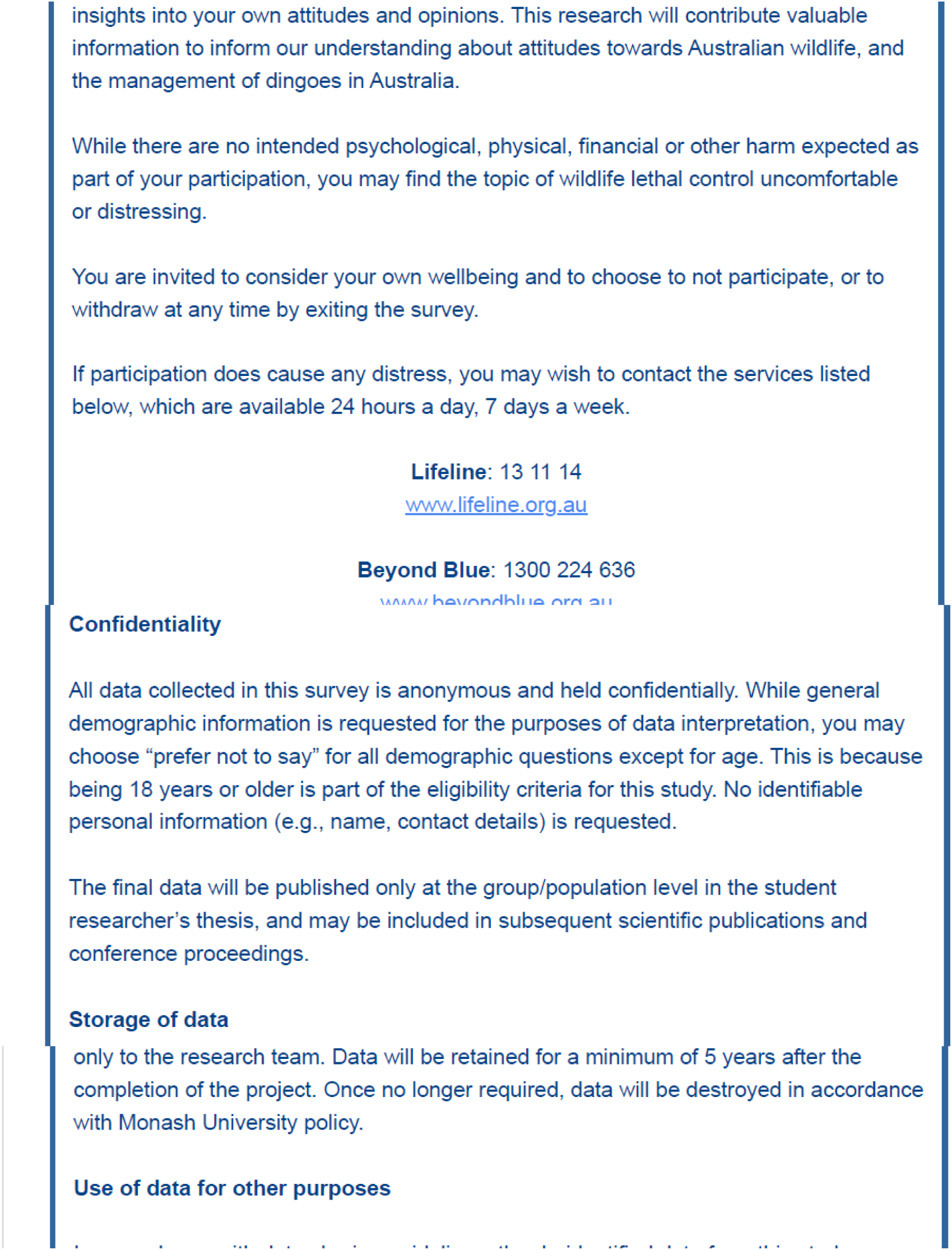
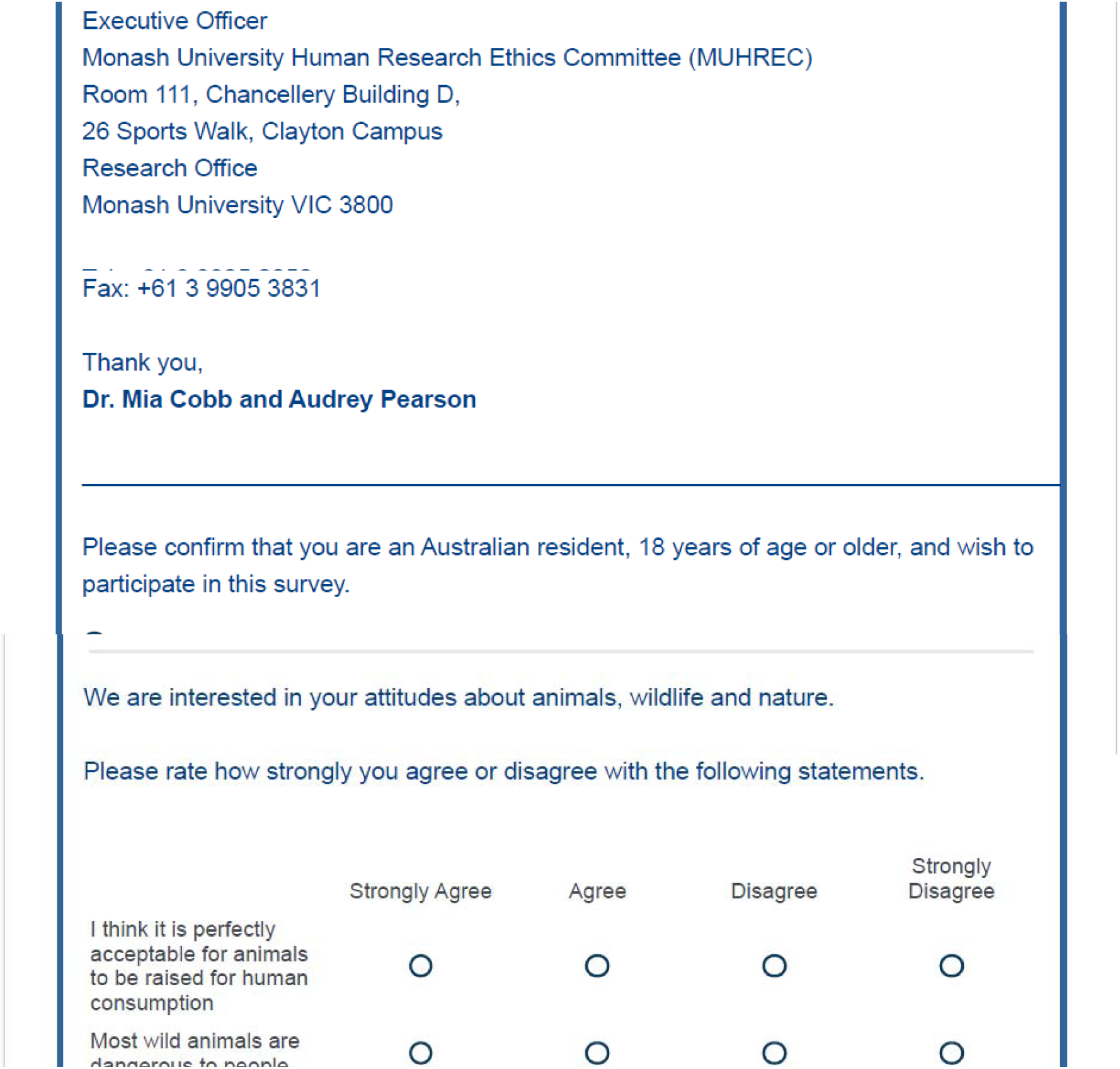
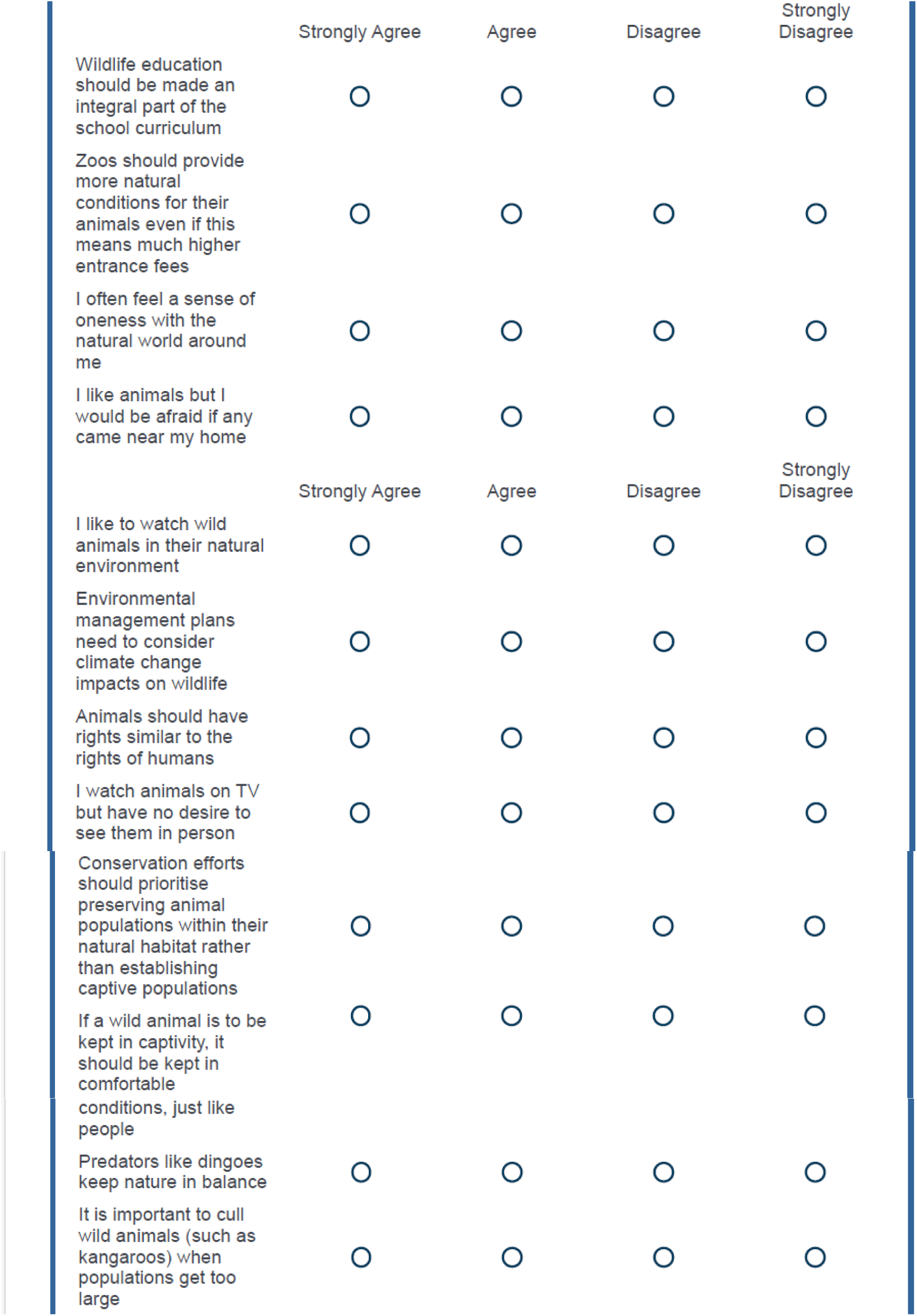
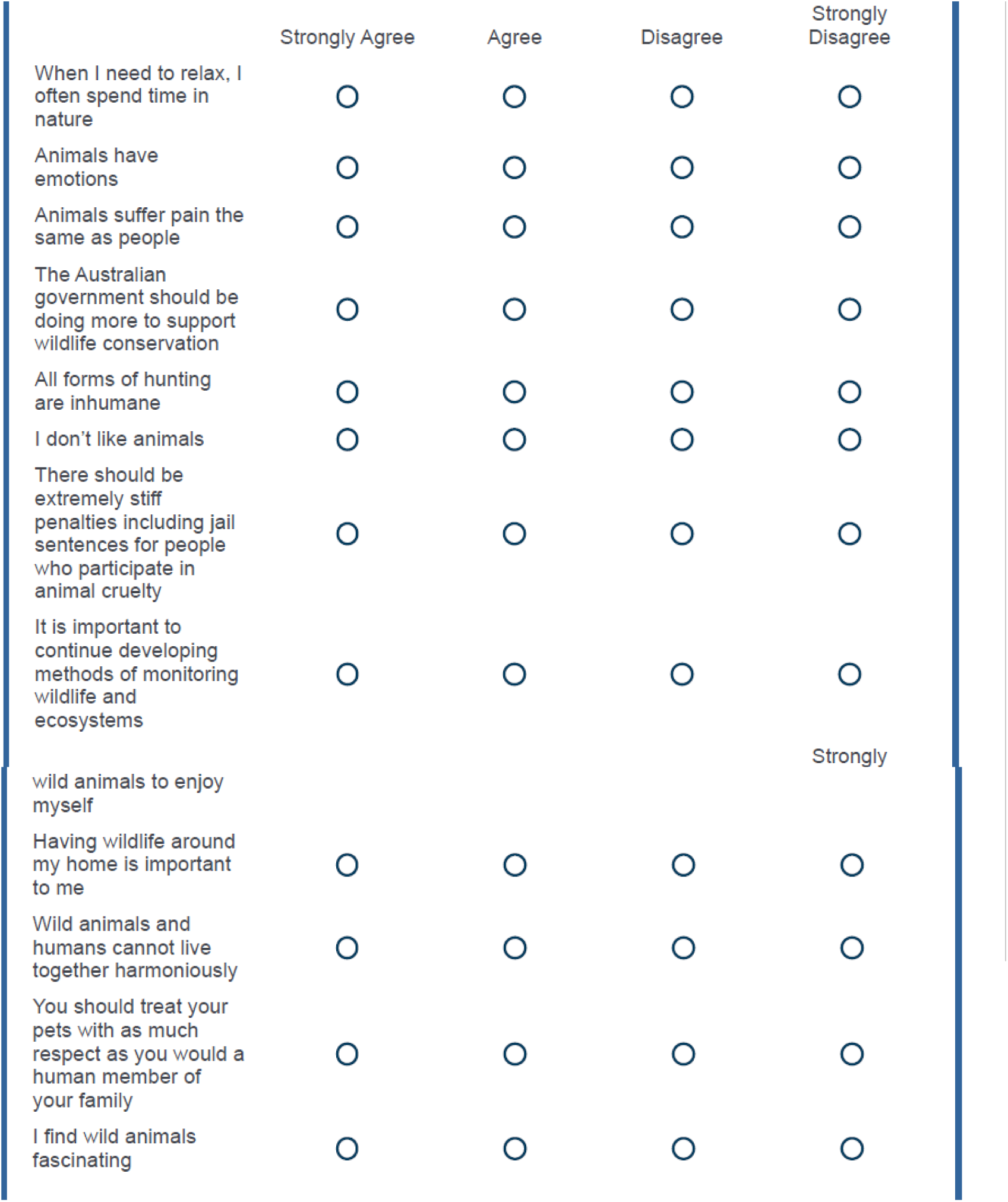
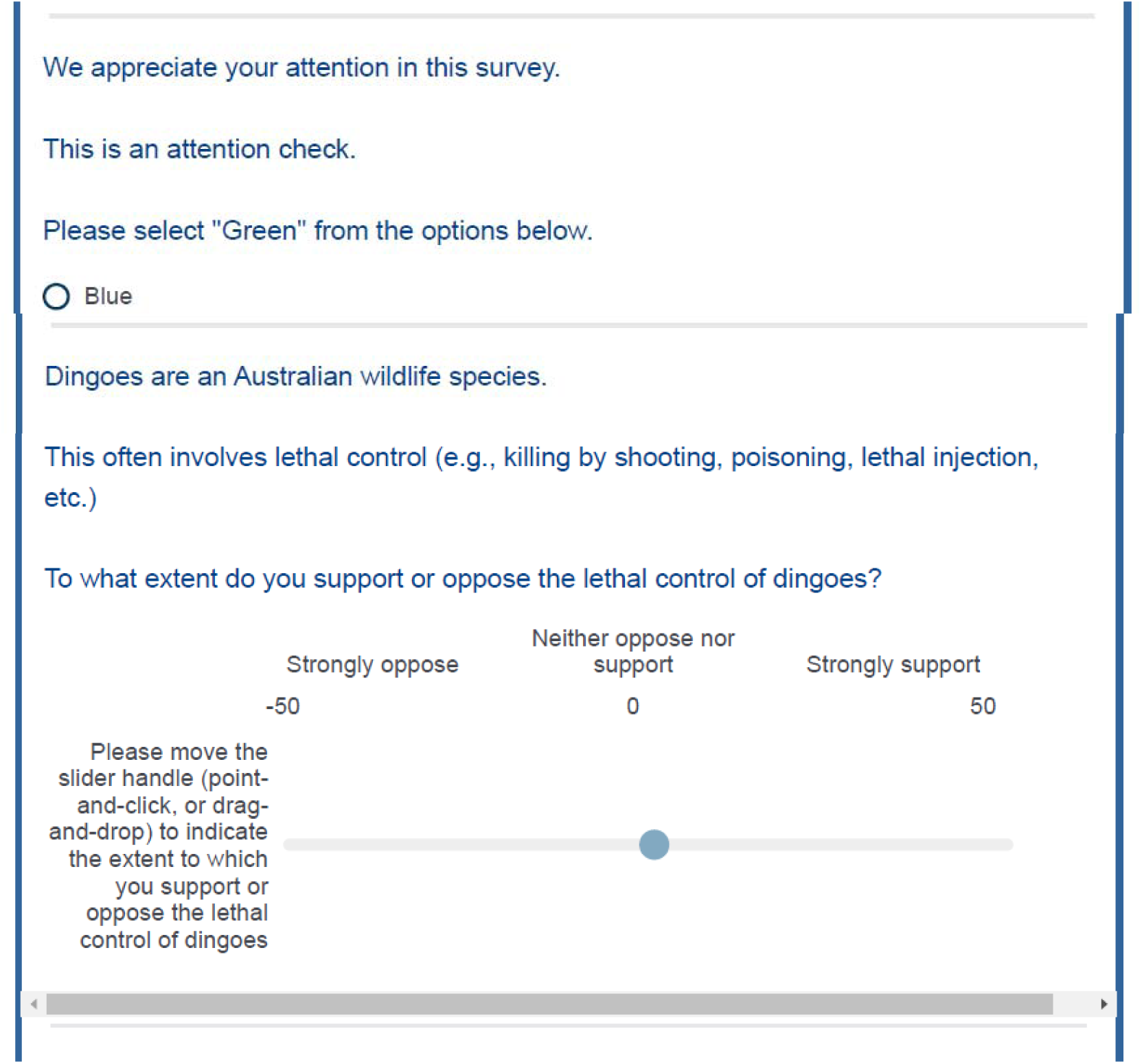
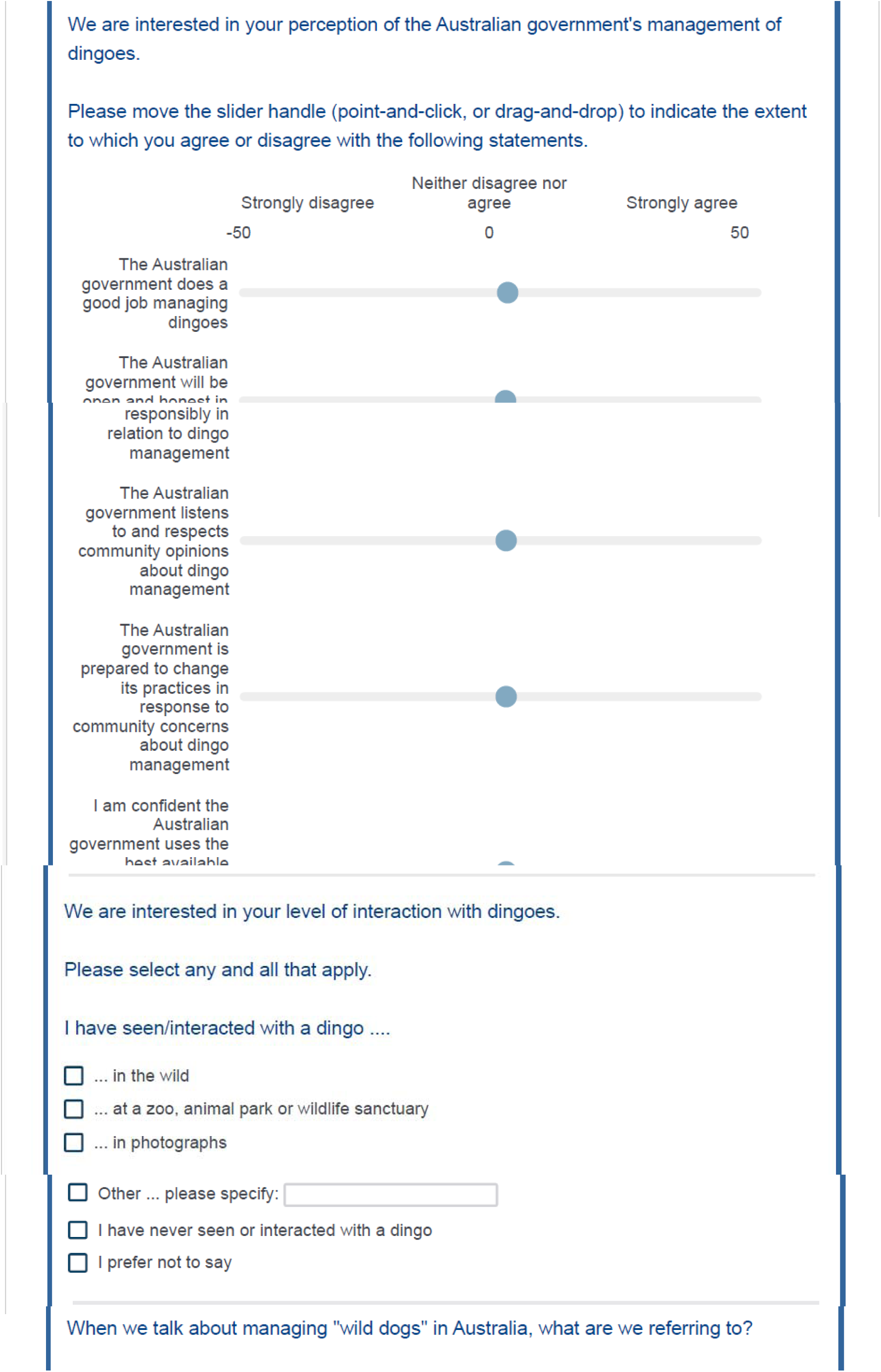
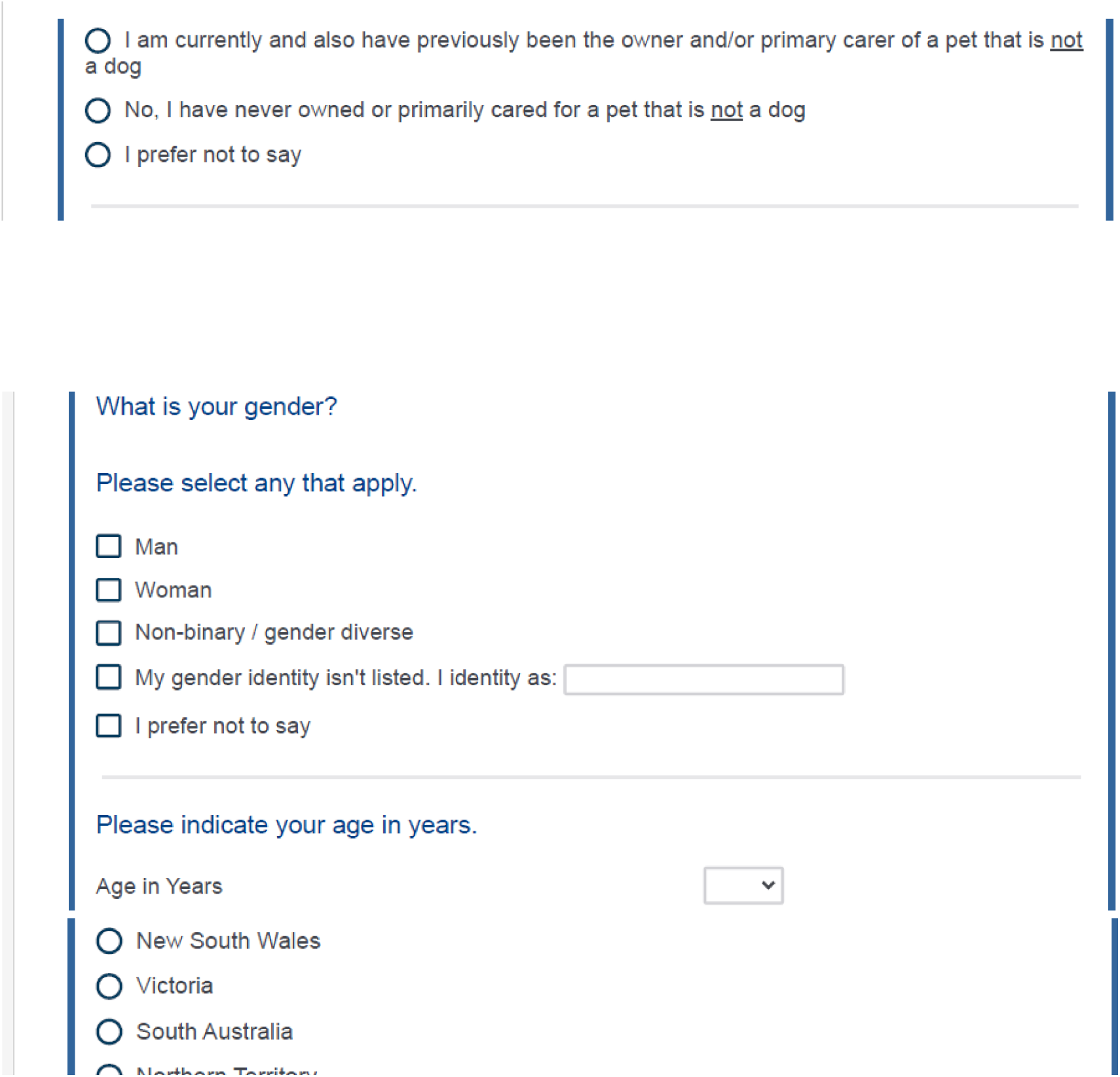
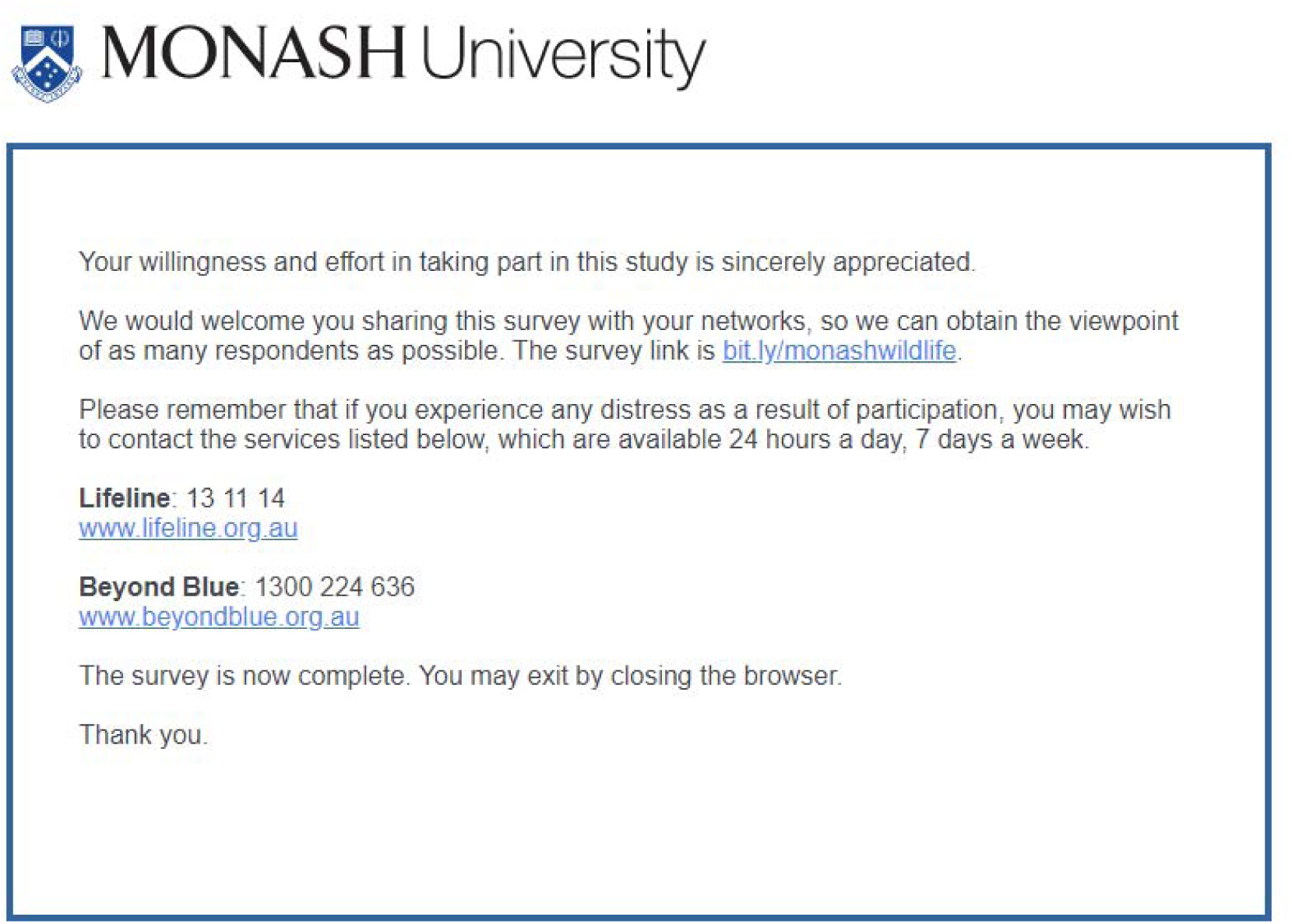

## Appendix C Data Cleaning and Assumption Testing

### Data Cleaning

Data cleaning was undertaken in Microsoft Excel (Office 365), and statistical analyses in IBM SPSS Statistics (Version 29). Open-text responses (i.e., “other – please specify”) for gender, education, and types of dingo interaction were recoded into existing or new categories to accurately reflect the demographic information. Items relating to social license pressure and support for lethal dingo control were recalculated to a 0 – 100 scale for analysis. Since incomplete responses were excluded, the analysed data set (*N* = 990) had no missing data. All scores were within expected ranges.

### Assumption Testing

#### One-Way Chi-Square Test

Prior to testing the hypothesis that more Australians would oppose (rather than support) lethal dingo control, the assumptions of a one-way chi-square test were examined following Witte and Witte’s (2015) recommendations. The assumption of independent observations was met with observed frequencies not exceeding the relevant category subtotals. Furthermore, all expected cell frequencies exceeded five. Therefore, all assumptions were met.

#### Mediation Model

The assumptions of a mediated regression model were examined before testing the predictive relationships between social license pressure, attitude towards Australian wildlife, and support for lethal dingo control, and whether the relationship between social license pressure and support for lethal dingo control was mediated by attitude towards Australian wildlife (H5). All model variables showed univariate non-normality with significant normality test results (*p* < .05), and standardised skewness (*Z_Skewness_*) and kurtosis (*Z_Kurtosis_*) statistics exceeding ±1.96 (*p* < .05, two-tailed), except kurtosis for social license pressure. See Table 1 for further detail.

Visual inspection of histograms and boxplots similarly indicated non-normality. Social license pressure was negatively skewed, with a peak of responses at the uppermost scale endpoint (high social license pressure). Attitudes to Australian wildlife was also negatively skewed with limited frequency counts observed at the lower end of the scale (negative attitudes). Support for lethal dingo control displayed a platykurtic, bimodal distribution with peaks at both endpoints (no support, full support), and a relatively flat distribution across all other scores. Following Field (2018), any potential bias from non-normality was considered sufficiently accounted for given the applicability of the central limit theorem (*N* > 30), the model being sufficiently powered (*N* > 105), and the use of bootstrapping as a robust estimation method.

Assumption testing for pathways *b c’*, *a*, and *c* was undertaken. Seven multivariate outliers were identified on pathway *b c’*, with Mahalanobis Distances’ greater than the critical value of 13.82 (*k* = 2, α =.001; Tabachnick & Fidell, 2019). Two of these multivariate outliers were also univariate outliers on the attitudes to Australian wildlife variable with standardised scores greater than ±3.29 (*p* < .001, two-tailed). Three further univariate outliers for pathway *b c’*, and one for pathway *a* were identified through casewise diagnostics with standardised residuals greater than ±3.29 (*p* < .001, two-tailed). However, maximum Cook’s Distance for all pathways was less than 1.00, which suggested there were no influential outliers. To confirm there was no outlier influence, the model outcome (pathway *ab*) was checked with, and without, all outliers. The model outcome did not change, therefore outliers were retained to improve the generalisability of results (Hair et al., 2014). The assumption of multicollinearity for pathway *b c’* was met (Field, 2018) as the bivariate Pearson’s correlation between predictor and mediator variable was .34 (less than .80), tolerance value was 0.88 (greater than 0.1), and variance inflation factor was 1.13 (less than 10).

Following Field’s (2018) guidelines, residuals on all pathways were assessed. Residual independence was indicated by Durbin-Watson statistics between 1.00 and 3.00. Using *zpred vs. zresid* scatterplots, linearity was indicated by a lack of curving, homoscedasticity by the constant variance of residuals across predicted outcome values, and normality by the even spread of points. Normality was also indicated if standardised residuals generally followed the normal distribution line on the Normal P-P Plot. All pathway residuals were linear, and independent with Durbin-Watson statistics of 1.79 (*b c’*), 1.55 (*a*), and 1.37 (*c*). While the Normal P-P Plots showed reasonable normality for pathway *b c’*, deviations were evident on pathways *a*, and *c*. The *zpred vs. zresid* scatterplots for all pathways displayed either inconsistent variance, or uneven spread across all predicted outcome values, suggestive of heteroscedasticity and non-normality. As such, the mediation model proceeded with bootstrapping (5,000 samples), and an HC3 Davidson-MacKinnon heteroscedastic-consistent approach to avoid biased standard errors and improve confidence in results (Allen et al., 2018; Tabachnick & Fidell, 2019).

#### Independent Samples T-Test

The assumptions of an independent samples *t*-test were examined before testing whether support for lethal dingo control significantly differed by dog ownership (dog-owned vs. dog-never-owned; participants who declined to answer were excluded). The assumptions of normality and homogeneity of variance were violated in the original model with uneven group sizes (dog-owned = 927; dog-never-owned = 56). Therefore, random sampling of the dog-owned group was undertaken to equalise group sizes at 56 participants. While independent samples *t*-tests do not require equal group sizes, confidence in results is improved when normality and homogeneity of variance assumptions are violated (Witte & Witte, 2015).

With equal group sizes, the dog-owned group exhibited a platykurtic, bimodal distribution with peaks at both endpoints (no support, full support), and a flat distribution across all other scores. The dog-never-owned group displayed a relatively normal distribution. Both groups violated normality tests (*p* < .05), however *Z_skewness_* only exceeded ±1.96 (*p* < .05, two-tailed) for the dog-owned group. No univariate outliers (standardised scores greater than ±3.29, *p* < .001, two-tailed) were identified for either group. The homogeneity of variance assumption was met with a non-significant Levene’s test (*p* = .074). Since *t*-tests are robust to small-moderate non-normality (Fein et al., 2022), and the overall sample size of 112 remained sufficiently powered, the analysis proceeded. See Table 2 for further detail.

## Footnotes

1 The study was funded by a government agency, and researchers were employed by the same funding source.

## Notes

### Competing Interest Statement

The authors have declared no competing interest.

